# A spatial model of the plant circadian clock reveals design principles for coordinated timing under noisy environments

**DOI:** 10.1101/2020.09.13.294785

**Authors:** Mark Greenwood, Isao T. Tokuda, James C.W. Locke

**Author notes:** **Abbreviations:** *CCA1*, *CIRCADIAN CLOCK ASSOCIATED 1*; CV, coefficient of variation; DD, constant dark; *ELF4*, *EARLY FLOWERING 4*; LD, light-dark; *LHY*, *LATE ELONGATED HYPOCOTYL*; LL, constant light; *LUC*, *LUCIFERASE*; *LUX*, *LUX ARRHYTHMO*; *PRR*, *PSEUDO-RESPONSE REGULATOR*; *TOC1*, *TIMING OF CAB EXPRESSION 1*; YFP, yellow fluorescent protein.

## Abstract

Individual plant cells possess a genetic network, the circadian clock, that times internal processes to the day-night cycle. Mathematical models of the clock network have driven a mechanistic understanding of the clock in plants. However, these models are typically either ‘whole plant’ models that ignore tissue or cell type specific clock behavior, or ‘phase only’ models that do not include clock network components explicitly. It is increasingly clear that in order to reveal the design principles of the plant circadian clock, clock network models must address spatial differences. This is because complex spatial behaviours have been observed in tissues and cells in plants, including period and phase differences between cells and spatial waves of gene expression between organs. Here, we implement an up to date clock network model on a spatial template of the plant. In our model, the sensitivity to light inputs varies across the plant, and cells communicate their clock timing locally via the levels of core clock mRNA levels by cell-to-cell coupling. We found that differences in sensitivities to environmental input in the model can explain the experimentally observed differences in clock periods in different organs, and we show using the model that a plausible coupling mechanism can generate the experimentally observed waves in clock gene expression across the plant. We then examined what features of the plant circadian system allow it to keep time under noisy light-dark (LD) cycles. We found that differences in sensitivity to light can allow regional flexibility in phase even under LD cycles, whilst local cell-to-cell coupling minimized variability in clock rhythms in neighboring cells. Thus, local sensitivity to environmental inputs combined with cell-to-cell coupling allows for flexible yet robust circadian timing under noisy environments.

## Introduction

The circadian clock is a 24 h genetic oscillator found in many organisms. The clock consists of a circuit of interlocking feedback loops of mRNAs and proteins that generate daily oscillations in circuit component levels. Signals from the environment align the timing of these oscillations to the day-night cycle [1]. Once set, circadian clocks act as an internal timing signal, allowing biological processes to anticipate the external environmental cycles. The clock modulates a diverse range of processes in plants, including cell division, tissue growth, flowering time, and scent emission [2–5]. Altogether this daily timing provides a significant fitness advantage to the plant [6,7].

Individual plant cells possess a robust circadian clock [8]. However, substantial differences in the period and phase of clock rhythms across the plant have been observed. Time lapse imaging experiments with luciferase and fluorescent reporter genes have shown that rhythms in core clock genes oscillate with different speeds in different organs under constant light (LL) [8–13]. Further, experiments under a range of conditions have shown that differences in clock speed and phase can be caused by organs having different sensitivity to environmental signals [12–15]. Differences in the clock network between tissues may also contribute to generating differences in rhythms across the plant, as although the clock genes are broadly expressed [13], some are tissue enriched [16], and mutations can affect organs differently [11,15,17].

The observed differences in clock rhythms across the plant raises the question of how clocks in different cells and tissues remain coordinated with each other. One mechanism would be for cells to communicate their timing with their neighbors. High resolution experiments have measured or inferred local coupling of clock rhythms between cells [8,11,14,16,18–20], and local coupling can drive spatial waves of clock gene expression across the plant [8,14,18–20]. Longer-distance coupling between clocks is also possible. For example, molecular signals communicate circadian temperature information from the shoot to the root [11,21], and light information may be piped down the stem to entrain the root [22].

Mathematical modeling has played a crucial role in gaining a mechanistic understanding of plant circadian clocks. Models of the network have increased in complexity over time in parallel with the growing number of experiments [23–28]. Recently, these detailed molecular models have been used to probe the differences between the shoot and root clock [13]. However, the coupling of clocks between cells was not considered. For this, more computationally tractable models of the clock network are necessary. Reduced models of the network have already been constructed that capture many of the features of the single cell clock dynamics [29–32]. However, these models have not been applied to study spatial dynamics. Instead, ‘phase only’ models that lack any genetic network information and only consider the phases of individual cellular rhythms have been preferred [8,18–20]. Although these models allow the simulation of general oscillatory behavior, owing to their simplicity they are unsuitable for investigating molecular mechanisms of the clock.

Recently, a ‘phase only’ Kuramoto model [33] was used to propose a mechanism for whole-plant coordination of clocks [8,14]. In order to match experimentally measured rhythms, the model fixed clock periods to different speeds in each region of the plant. It assumed faster rhythms in the cotyledons, hypocotyl, and root tip, and slower rhythms in the rest of the root, as observed experimentally. With these periods fixed, cells were allowed to communicate clock phase through local cell-to-cell coupling. With these assumptions, simulations of the model generated waves of clock gene expression within and between organs, as observed experimentally [8,14]. Thus, local cell-to-cell coupling could enable coordination between organs in plants.

Multiple questions remain about how the plant clock coordinates rhythms that cannot be addressed using a ‘phase only’ model. For example, how are the periods set differently in different parts of the plant? What molecular mechanisms allow the coupling of clock rhythms from cell-to-cell? How can the plant clock network ‘filter’ both internal and environmental noise to robustly entrain to the environment? To begin to address these questions, in this work we developed a spatial network model of the plant circadian system. We modified a previously generated simplified network model and implemented it on a multicellular template of a plant. In our model, the sensitivity to light varies across the plant, and cells communicate via the mRNA levels of a core clock gene. Simulations capture the organ specific period differences and spatial waves observed under LL, demonstrating a plausible mechanism of circadian coordination. We then applied our model to examine how the system keeps time under noisy light-dark (LD) cycles. We found that regional differences persist even under LD cycles, but cell-to-cell minimized differences between neighbor cells. Thus, the combination of local sensitivity to inputs and cell-to-cell coupling allows for coordinated timing in noisy environments.

## Results

### A locally coupled spatial model of the plant circadian clock network

We first implemented a reduced network model of the *Arabidopsis thaliana* circadian clock [31]. To decrease the complexity of the model, the authors grouped functionally similar genes into single entities (Fig 1A). The compact network model incorporates known light inputs to the network and qualitatively recapitulates clock dynamics under both light dark cycles and constant light. At only 9 equations and 34 parameters the model is also computationally tractable for spatial simulations. We modified the De Caluwe model to include a repression rather than activation interaction between *PSEUDO-RESPONSE REGULATOR 9 (PRR9*)/*PRR7* and *CIRCADIAN CLOCK ASSOCIATED 1* (*CCA1*) / *LATE ELONGATED HYPOCOTYL* (*LHY*), as this has recently been shown experimentally [34] (Methods).

**Fig 1.**
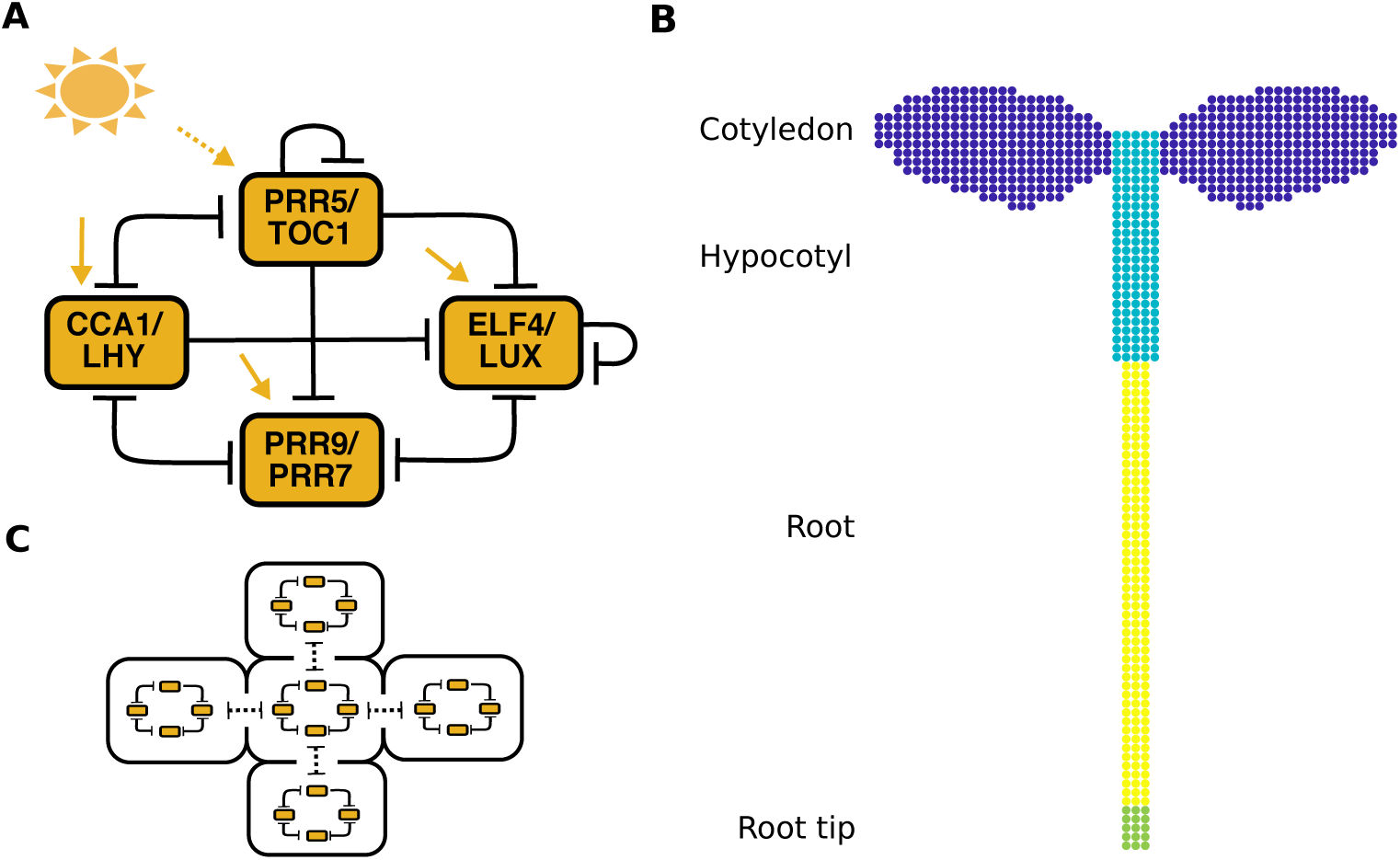
The structure of the spatial circadian clock model. (A) Summary of the modified compact circadian clock model used for simulations. The original compact model [31] is modified to include a repression interaction between CCA1/LHY and PRR5/TOC1. Light input is received by all gene pairs, but affects their production and degradation rates to different degrees. Yellow dashed arrows represent light input at the transcriptional level and solid yellow arrows represent light input at both the transcriptional and post-transcription level. “T” arrows represent repression. (B) The network was implemented within each cell on a simplified template of a seedling, with cells classified as either cotyledon (blue), hypocotyl (turquoise), root (yellow), or root tip cells (green). (C) The network in each individual cell is coupled to the neighboring cells through the level of *CCA1*/*LHY*. *CCA1*, *CIRCADIAN CLOCK ASSOCIATED 1*; *ELF4*, *EARLY FLOWERING 4*; *LHY*, *LATE ELONGATED HYPOCOTYL*; LL, constant light; *LUC*, *LUCIFERASE*; *LUX*, *LUX ARRHYTHMO*; *PRR*, *PSEUDO-RESPONSE REGULATOR*; *TOC1*, *TIMING OF CAB EXPRESSION 1*.

We implemented the modified De Caluwe model on a simplified template of a seedling. The template consisted of approximately 800 cells, classified into cotyledon, hypocotyl, root, and root tip regions (Fig 1B and Methods). Although a number of studies have demonstrated local cell-to-cell coupling between clocks in *A. thaliana* [8,11,14,16,18–20], the identity of the coupling components are unclear. Initially, to model the coupling (Fig 1C), the level of *CCA1*/*LHY* in one cell was assumed to be locally coupled to the level of the cell’s neighbors. The coupling strength was set to 2 (Methods). To simulate the variability observed in single cell clock rhythms [8] we multiplied the level of each mRNA and protein by a time scaling parameter that was randomly selected from a normal distribution (Methods). The range of this normal distribution was set differently for each organ to qualitatively match the variability observed experimentally (S1 Fig).

### Different light sensitivities can explain organ differences in phase and period

We next attempted to recapitulate in our model the differences in clock period and phase in different organs that have been observed in experiments using a single cell CCA1-YFP reporter [8] and a *GIGANTEA* luciferase reporter [14]. In these experiments, faster rhythms were observed in the cotyledon, hypocotyl, and root tip, with slower rhythms in the rest of the root. We first reanalyzed existing luciferase data [14] and confirmed that these relationships held for several of the core clock genes in our model, *PSEUDO-RESPONSE REGULATOR 9* (*PRR9)*, *TIMING OF CAB EXPRESSION 1* (*TOC1*), and *EARLY FLOWERING 4* (*ELF4*) (Fig 2A). Whereas in our previous ‘phase only’ model, we fixed the periods to be different in each part of the plant, with our spatial network model, we could now investigate what causes the differences in periods. Previously it has been hypothesized that different sensitivities to environmental inputs alter periods across the plant [13,14]. For example, cells in the cotyledon may be fast because they are more sensitive to light. Experimental tests of this, however, are confounded by changes in metabolism and development caused by light [35]. To test this in our model, we simulated differences in sensitivity by setting the intensity of light input in our model to differ depending on the region, and examined whether this can generate the period differences observed across the plant.

**Fig 2.**
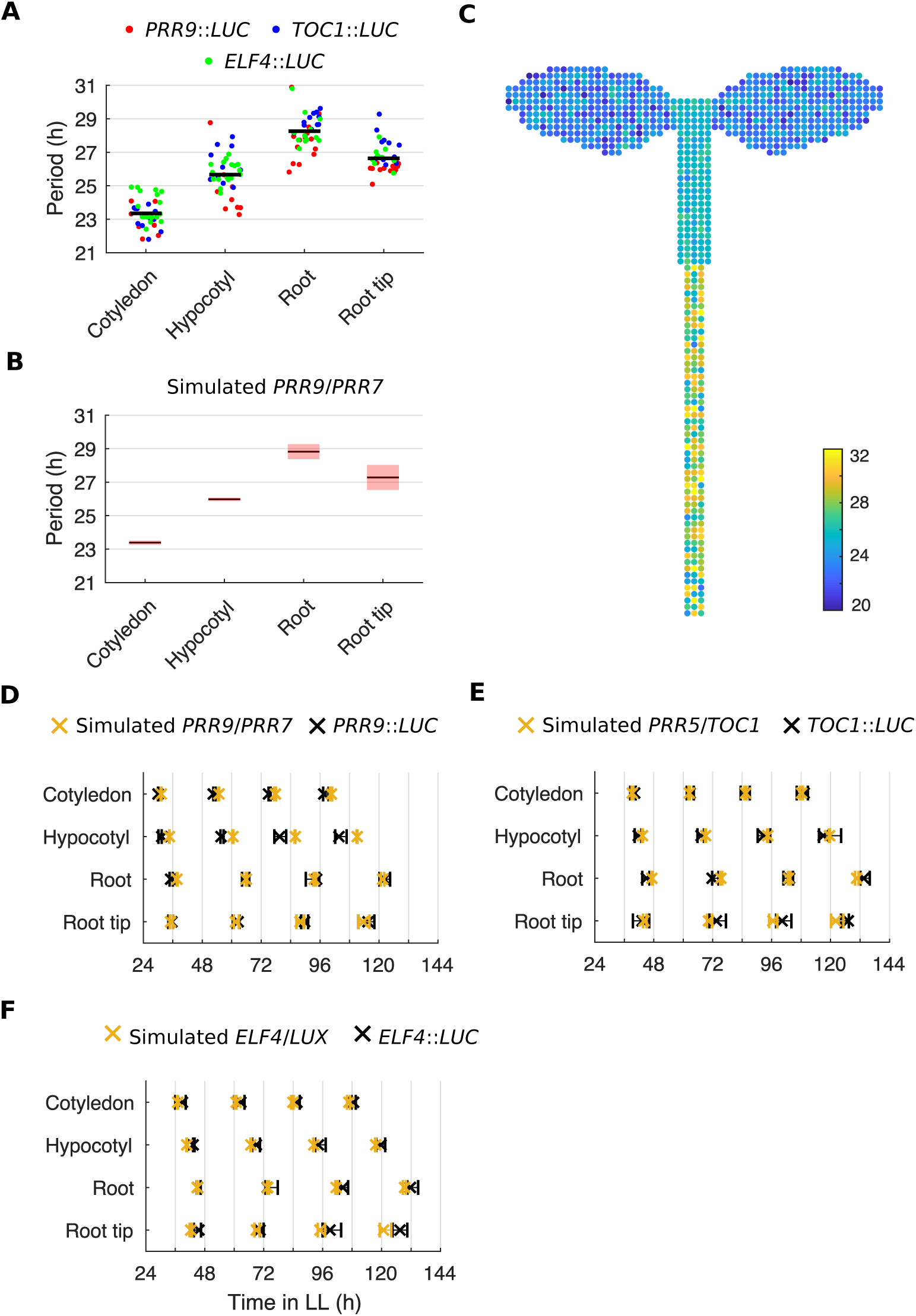
Regional differences in light input can generate the period structure observed experimentally. (A) Period estimates of *PRR9*::*LUC*, *TOC1*::*LUC*, and *ELF4*::*LUC* for different organs imaged under LL. Each data point represents a period estimate from the organ of a single seedling. The horizontal black line shows the mean. (B) Period estimates of simulated *PRR9/PRR7* expression, measured from regions within the template seedling. The black line indicates the mean and the red shaded area one standard deviation of 18 independent simulations. (C) Periods of individual cells plotted on the template seedling. The color of the cell represents the speed of the oscillation. By assuming higher light input to the cells in the cotyledon, hypocotyl, and root tip, the model approximates the period differences observed between regions in experiments, A. A noise parameter was adjusted for each region to simulate the variability of periods observed in single cell experiments (S1 Fig). (D–F) Times of peaks of simulated *PRR9*/*PRR7* and *PRR9*::*LUC* (D), simulated *PRR5*/*TOC1* and *TOC1*::*LUC* (E), or simulated *ELF4*/*LUX* and *ELF4*::*LUC* (F), in different organs under LL. Simulations include different levels of light input to regions and cell-to-cell coupling (*J* = 2). Plots represent the 25th percentile, median, and the 75th percentile for the peak times of oscillations scored as rhythmic, *n* = 9 simulations. Experimental data is an analysis of time-lapse movies carried out previously [14]. *CCA1*, *CIRCADIAN CLOCK ASSOCIATED 1*; CV, coefficient of variation; *ELF4*, *EARLY FLOWERING 4*; *LHY*, *LATE ELONGATED HYPOCOTYL*; LL, constant light; *LUC*, *LUCIFERASE*; *LUX*, *LUX ARRHYTHMO*; *PRR*, *PSEUDO-RESPONSE REGULATOR*; *TOC1*, *TIMING OF CAB EXPRESSION 1*.

We entrained the cells in our simulations to LD cycles for 4 days before releasing them into LL for a further 6 days and measured the periods, as carried out in previous experiments [14]. When assuming high sensitivity to light in the cotyledon (*L* = 1.35) and hypocotyl (*L* = 1.10), but lower in the root (*L* = 0.90) and root tip (*L* = 1.03), all organs entrained to the LD cycles (S2 Fig). Upon transfer to LL, we were able to generate different periods (Fig 2B) and phases (Fig 2D–F and S3 Fig) in each organ, matching those observed experimentally. Thus, our results revealed that different sensitivities to environmental inputs are sufficient to generate the experimentally observed spatial differences in period and phase across the plant. This is due to higher light sensitivity causing the clock to run faster in our simulations (S4 Fig), as expected for a diurnal organism [36].

### Local sharing of clock mRNA levels can drive spatial waves of clock gene expression

Previously we observed two waves of clock gene expression, one traveling up, and one down the root, in *CCA1*, *PRR9* and *GI* reporters [8,14]. These waves could be explained in a phase only model by local cell-to-cell coupling [8,14,19,20]. We next tested whether a plausible mechanism for cell-to-cell coupling, sharing of mRNA between cells [37], can recapitulate the experimental observations. To provide a benchmark, we first analyzed the seedlings carrying transcriptional reporters for *TOC1*, *PRR9*, and *ELF4* [14] at the sub-tissue level (Methods). As in previous studies [8,14,18–20], space-time plots revealed spatial waves of gene expression within and between organs (Fig 3A and S5 Fig). The direction of the waves can be clearly observed in plots of the final peaks of expression (Fig 3B and S5 Fig). For each gene, the wave patterns appeared similar, with two waves of clock gene expression in the root (Fig 3C, S5 Fig, and S1 Video).

**Fig 3.**
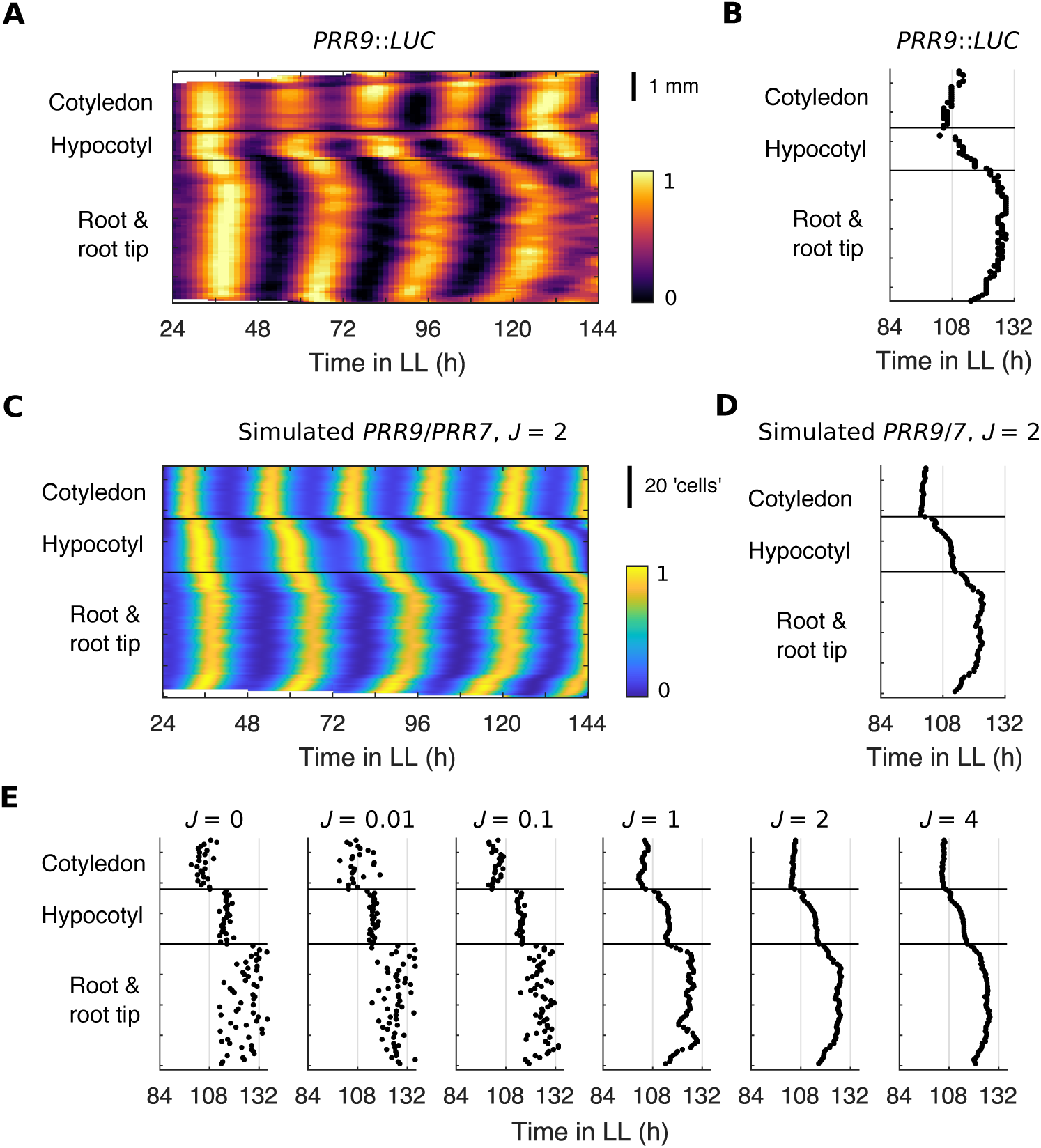
Local sharing of mRNA can reproduce experimentally observed spatial waves of clock gene expression. (A) Representative intensity plot of *PRR9*::*LUC* expression across longitudinal sections of a single seedling under LL. (B) Times of the final peaks of the *PRR9*::*LUC* intensity plot simulated under LL. (C) Representative intensity plot of simulated *PRR9*/*PRR7* expression across longitudinal sections of a single seedling under LL. Simulations include different levels of light input to regions and cell-to-cell coupling (*J* = 2). (D) Times of the final peaks of the simulated *PRR9*/*PRR7* intensity plot simulated under LL. (E) Times of the final peak of simulated *PRR9*/*PRR7* intensity plots, each simulated under LL with increasing strength of coupling, *J*. Experimental data is an analysis of time-lapse movies carried out previously [14]. *PRR*, *PSEUDO RESPONSE REGULATOR*; LL, constant light; *LUC*, *LUCIFERASE*. *TOC1, TIMING OF CAB EXPRESSION 1*.

We next analyzed simulations at the sub-tissue level (S6 Fig and Methods) to see if our model captures these spatial dynamics. For each gene, the wave patterns appeared similar to experiments, traveling from the fast oscillating regions that are more sensitive to light, into the slower regions with lower sensitivity to light (Fig 3C, D, S5 Fig and S2 Video). These waves required cell-to-cell coupling through the local sharing of clock mRNA, as we only observed waves with coupling strengths, *J*, above approximately 1 (Fig 3E and S7 Fig). Similar simulation results were obtained using different clock genes as the coupling component, suggesting that any cell-to-cell sharing of clock components can explain the experimentally observed spatial dynamics (S8 Fig). We also found that our simulation results were qualitatively similar when keeping regional differences of light sensitivity and local cell-to-cell coupling, but assuming no cell-to-cell variability within regions (S9 Fig). Additionally, we ran simulations assuming longer distance and global coupling. Increasing the local coupling to be between 8 rather than 4 neighbor cells gave qualitatively similar results. However, with global coupling all cells adopted the same phase at higher coupling strengths, regardless of position in the plant (S10 Fig). Finally, by setting the light inputs in our locally coupled model to be equal in all regions of the seedling, we were also able to simulate the loss of waves observed in the light sensing mutant *phyb-9* [14] (S11 Fig). Taken together, these results show that the assumptions of local cell-to-cell coupling and differential light sensitivity between regions are the key aspects of our model that allow a match to experimental data.

### Local flexibility persists under idealized and noisy LD cycles

Our modeling suggests that different sensitivities to environmental inputs across the plant allows the clock to be locally flexible under LL, by allowing it to adopt different periods and phases across the plant even under the same environment. We simulated our model under LD cycles to investigate whether local flexibility persisted with rhythmic input. We first simulated idealized LD cycles, where lights are switched fully on during the daytime and off at night (Fig 4A and Methods). The sensitivity to light during the daytime differed depending on the region, as with LL simulations, and we first assumed no cell-to-cell coupling (*J* = 0). In the absence of cell-to-cell coupling, close inspection of the final peaks of expression revealed small differences in the timing between regions. *PRR5*/*TOC1* expression in cells of the cotyledon peaked at 132.70 ± 0.50 h (median ± interquartile range), followed by those in the hypocotyl (133.40 ± 0.40 h) and then the root, with cells at the tip (133.50 ± 1.35 h) peaking before those in the middle region of the root (134.80 ± 1.20 h; Fig 4B, left and S12 Fig). These phase differences were qualitatively the same as observed under LL (Fig 2 and 3) but were discontinuous between regions. With the addition of cell-to-cell coupling, the phase differences between regions were no longer discontinuous and were instead staggered, causing the emergence of spatial waves (Fig 4B, middle and right, and S12 Fig). However, phase differences between the regions persisted and did not reduce in size with increasing strengths of coupling (S12 Fig). For example, there was a difference between the cotyledon and root of 1.90 ± 0.18 h (median ± interquartile range) without coupling (*J* = 0), 2.00 ± 0.20 h with coupling (*J* = 2), and 2.00 ± 0.18 h with stronger coupling (*J* = 4). A similar spatial structure was observed in *CCA1*/*LHY* and *PRR9*/*PRR7* expression (S12 Fig). However, *ELF4*/*LUX* appeared more synchronized when the peak coincided with the dark transition, as darkness causes strong repression of *ELF4* expression.

**Fig 4.**
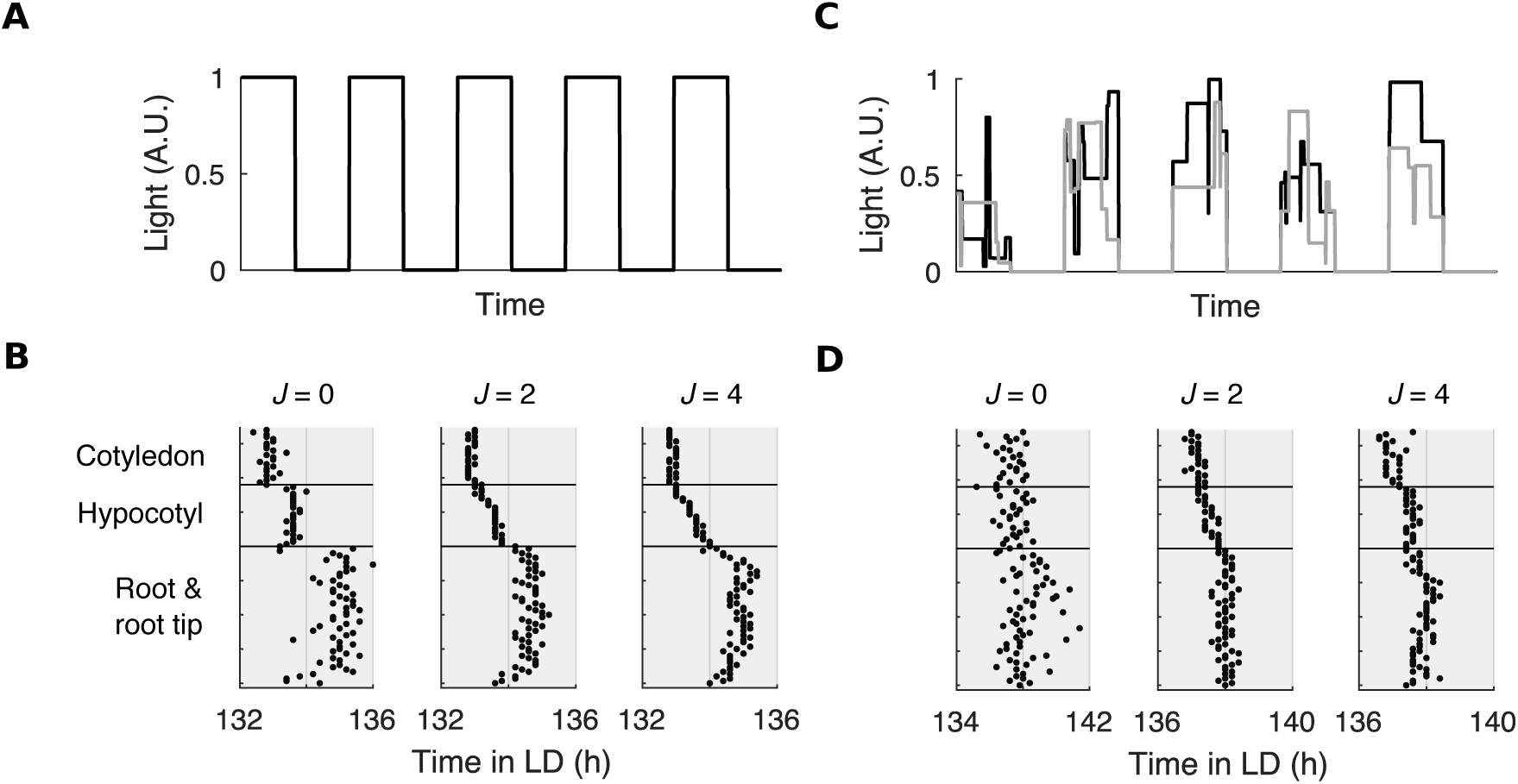
Regional phase differences persist under idealized and noisy LD cycles. (A) Schematic of idealized LD cycles, without fluctuations in light levels during the daytime. (B) Times of the final peak of simulated *PRR5*/*TOC1* intensity plots, each simulated under idealized LD with increasing strength of coupling, *J*. (C) Schematic of noisy LD cycles, with fluctuations in light levels during the daytime. Each line represents an LD cycle for a single cell. (D) Times of the final peak of simulated *PRR5*/*TOC1* intensity plots, each simulated under noisy LD with increasing strength of coupling, *J*. LD, light-dark; *PRR*, *PSEUDO-RESPONSE REGULATOR*; *TOC1*, *TIMING OF CAB EXPRESSION 1*.

We next simulated more realistic LD cycles containing fluctuations in the light intensity (Fig 4C). These LD cycles were designed to approximate the differences in input that cells may experience due to weather or a cell’s microenvironment (Methods). Examination of the times of the final peaks of expression reveals large variation in peak times within regions when cell-to-cell coupling was absent (*J* = 0; Fig 4D, left). This made any difference in phase between regions difficult to resolve. However, with increasing strengths of cell-to-cell coupling we observed a decrease in variability. For example, there was a difference of 5.00 ± 1.05 h (median ± interquartile range) between the earliest and latest peaking cells without coupling (*J* = 0), but 3.10 ± 0.23 h with coupling (*J* = 2), and 3.20 ± 0.15 h with stronger coupling (*J* = 4). This decrease in cell-to-cell variation revealed an underlying spatial structure (Fig 4D, middle and right, and S13 Fig), comparable to that observed under idealized LD cycles (Fig 4B, middle and right, and S12 Fig). Phase differences and spatial waves persisted with increasing strengths of coupling (S13 Fig). This spatial structure was also observed in expression of *CCA1*/*LHY* and *PRR9*/*PRR7*, but not in *ELF4*/LUX expression (S13 Fig). Together this shows that local flexibility in phase, and spatial waves, persisted under LD cycles when local cell-to-cell coupling was present.

### Cell-to-cell coupling maintains global coordination under noisy light-dark cycles

Simulations under noisy LD cycles showed that whilst local flexibility of regions persisted, cell-to-cell coupling decreased the phase variation within regions (Fig 4). We further investigated the function of cell-to-cell coupling by testing whether coupling also affected the global synchronization of rhythms, or is limited to a local effect on rhythms. To test this, we quantified the synchrony between all cells within the template seedling using a score for synchronization, *R* [38]. The synchronization index, *R*, is the ratio of the variance of the average signal to the average variance of the individual cells. When cells are synchronized these variances are equal to each other and *R* becomes 1, whereas when desynchronized *R* becomes 0. For each gene pair, we observed a high *R* value under idealized LD cycles, with or without cell-to-cell coupling (Fig 5, black lines). However, cells were less synchronous under noisy LD cycles (Fig 5, red lines). Each gene showed a reduced level of synchrony in the absence of cell-to-cell coupling (*J* = 0). *PRR9*/*PRR7* was the most affected by the noise, with a synchronization index of 0.97 ± 0.002 (mean ± standard deviation) under idealized LD and 0.61 ± 0.008 under noisy LD cycles (Fig 5B). This is likely because *PRR9*/*PRR7* synthesis is most strongly affected by light (Table 1).

**Fig 5.**
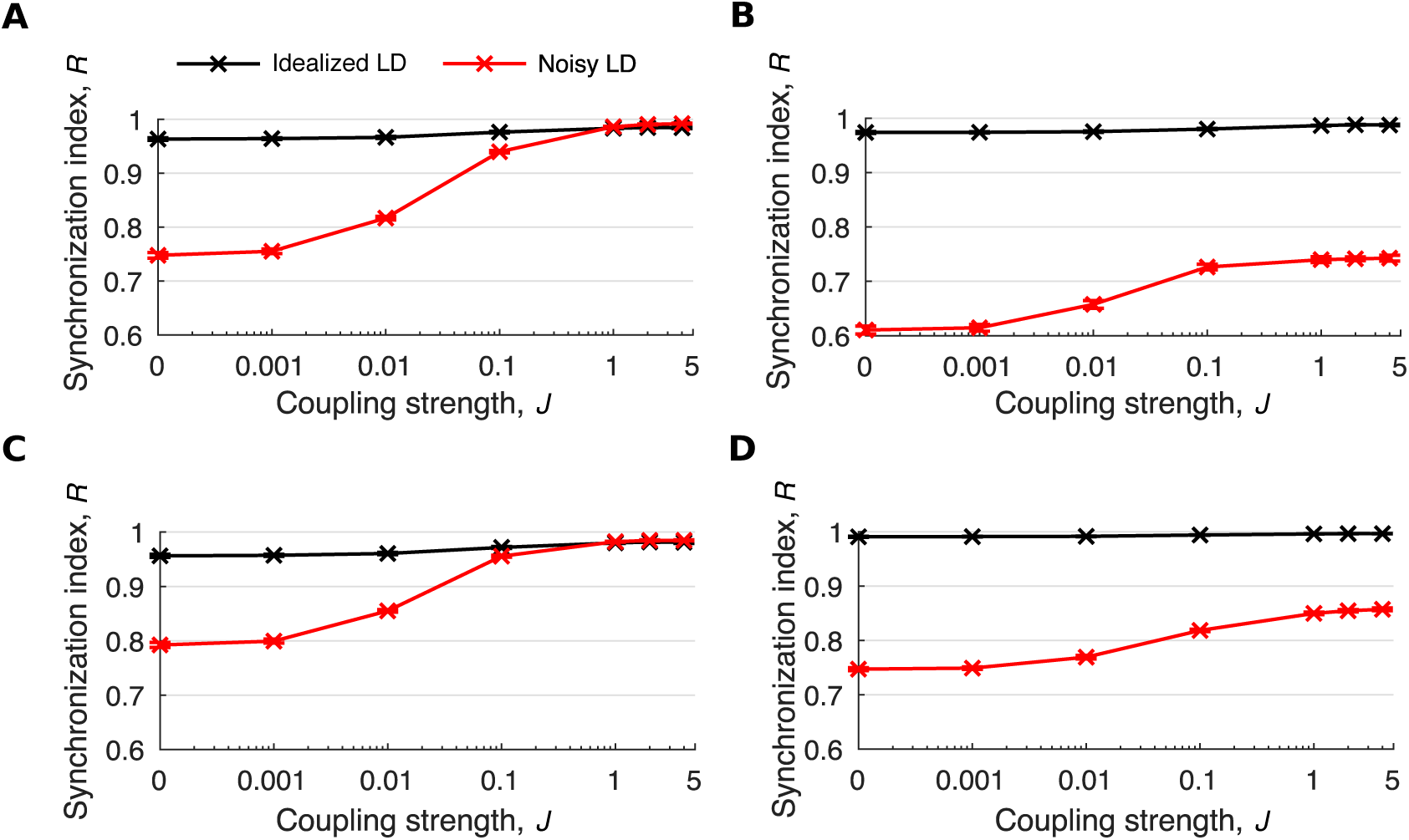
Cell-to-cell coupling maintains global synchrony under noisy LD cycles. (A–D) Quantification of phase coherence by the synchronization index, *R*, for simulated *CCA1*/*LHY* (A), *PRR9*/*PRR7* (B), *PRR5*/*TOC1* (C) or *ELF4*/*LUX* (D) mRNA expression under idealized or noisy LD cycles. Simulations included increasing strengths of coupling, *J*. Color legends are as in A. Data points represent the mean ± standard deviation, *n* = 9 simulations. *CCA1*, *CIRCADIAN CLOCK ASSOCIATED 1*; *ELF4*, *EARLY FLOWERING 4*; LD, light-dark; *LHY*, *LATE ELONGATED HYPOCOTYL*; *LUX*, *LUX ARRHYTHMO*; *LUC*, *LUCIFERASE*; *PRR*, *PSEUDO-RESPONSE REGULATOR*; *TOC1*, *TIMING OF CAB EXPRESSION 1*.

We observed an increase in global synchronization under noisy LD cycles as coupling strengths increased (Fig 5). At higher strengths of coupling (*J* > 2), for *CCA1/LHY* and *PRR5/TOC1* the synchronization index under noisy LD cycles approximately equaled that under idealized LD cycles (Fig 5A and C). For *PRR9*/*PRR7* and *ELF4*/*LUX* expression, the synchronization index increased with increasing strength of coupling, but did not reach the level observed under idealized LD cycles for all strengths tested (Fig 5B and D). Oscillations of protein expression also increased in synchrony with cell-to-cell coupling (S14 Fig). Thus, local cell-to-cell coupling improves the global coordination of circadian timing under noisy light-dark cycles, whilst still allowing differences in clock timing between regions.

## Discussion

Here we develop a spatial network model for the plant clock and use it to examine the design principles of clock coordination in plants. This simple multicellular model, which assumes regional differences in sensitivity to light and cell-to-cell coupling through the local sharing of clock mRNA levels, successfully simulates the period differences and phase waves across the plant observed under LL in previous time lapse experiments. Our simulations predict that phase flexibility persists under LD cycles, with cell-to-cell coupling reducing cell-to-cell differences but still allowing regional flexibility. We therefore find that the plant circadian clock system can combine regional differences in environmental signaling with cell-to-cell coupling to enable robust, yet flexible, circadian timing under noisy environmental cycles.

Previous models of the plant circadian clock have proven important for understanding aspects of plant physiology, including starch metabolism and flowering [39,40]. However, it was recently shown that the clock controls physiology in a tissue-specific manner. For example, clocks in the epidermis regulate growth, whereas those in the vasculature regulate flowering [41]. In future, our spatial model could be extended to investigate circadian control of physiology at the cell and tissue level. An interesting example will be cell division, which is regulated by the clock [2], and occurs in meristematic regions of the plant [42]. It will be interesting to see if, and how, flexibility of the circadian system impacts tightly controlled phenotypes such as this.

Including noise in LD cycles represents a step towards understanding how the plant circadian system works under natural conditions. Our simulations under noisy LD cycles reveal that local cell-to-cell coupling allows a balance between ensuring cells are coordinated locally even under a noisy environment, whilst still allowing regional differences that would not be possible with a global coupling mechanism. However, this is still a simplification of the real environment. For example, we do not consider temperature, which is phase shifted between the air and soil [43]. Although technically challenging, experiments under more realistic conditions such as this will be important for furthering our understanding. Promising advances in this direction include the GLO-Roots system [44], and the robotics of Bordage et al. [13], which allow the root to be imaged in the dark under LD cycles. As experiments progress, we predict that spatial models such as ours will be crucial for understanding how the clock integrates such a range of environmental signals across the plant.

The molecular details underlying the spatial differences, and coordination, of the plant circadian system are still being deciphered. Recently, *ELF3*, *ELF4*, and *LUX* were found to be central to the difference between the shoot and root, possibly due to their interaction with the light signaling protein PHYB [15]. However, interpretation is hindered by the fact that components causing differences could also act as a cell-to-cell mobile coupling signal. For example, ELF4 has also been proposed to move from the shoot to couple with clocks in the root [21]. We note that our spatial molecular model has the potential to delineate these two effects. In future work it will be important to vary the identity of the coupling signal, and the underlying genetic network, to generate testable predictions.

The plant circadian system, with local inputs to cells that are coupled together, represents a decentralized structure. This is in contrast to the centralized mammalian circadian system [45]. Here we found that this decentralized structure can afford plants flexibility, allowing regions to adopt different phases under noisy LD cycles. It could be speculated that this has a physiological effect on plants, causing, for example, a stagger in the timing of growth across the plant. However, further work is required to understand how, or whether, this flexibility is an advantage to plants.

## Acknowledgments

MG and JL were supported by the Gatsby Charitable Foundation (grant number GAT3395/GLC) and IT by the Japan Society for the Promotion of Science (grant numbers 17H06313, 18H02477, and 20K11875). We thank Dr. Katie Abley (University of Cambridge) for critical reading of the manuscript.

## Methods

### Single cell molecular model

As a model for the plant circadian clock, we exploit the compact model introduced by De Caluwe et al. [31]. The original compact model consists of 9 ordinary differential equations. Among them, 8 equations describe the temporal evolution of the mRNA and protein levels of the core clock genes. The clock genes are grouped into four sets of lumped pairs labeled as: CL (*CCA1* and *LHY*), P97 (*PRR9* and *PRR7*), P51 (*PRR5* and *TOC1*) and EL (*ELF4* and *LUX*). The 9-th equation is for the light-sensitive protein P (PIF3 and PIL1).

In our modeling, one modification to the original compact model was made. According to the experiments reported in [46], the compact model assumed that the P97 variable is activated by CL. However, more recent work [28,34], has shown that *LHY* acts as a repressor of all other clock components, including *PRR9* and *PRR7*. We therefore replaced the activation term that represents the connection from CL to P97 with a repression. The revised single cell model is now described by the following differential equations:

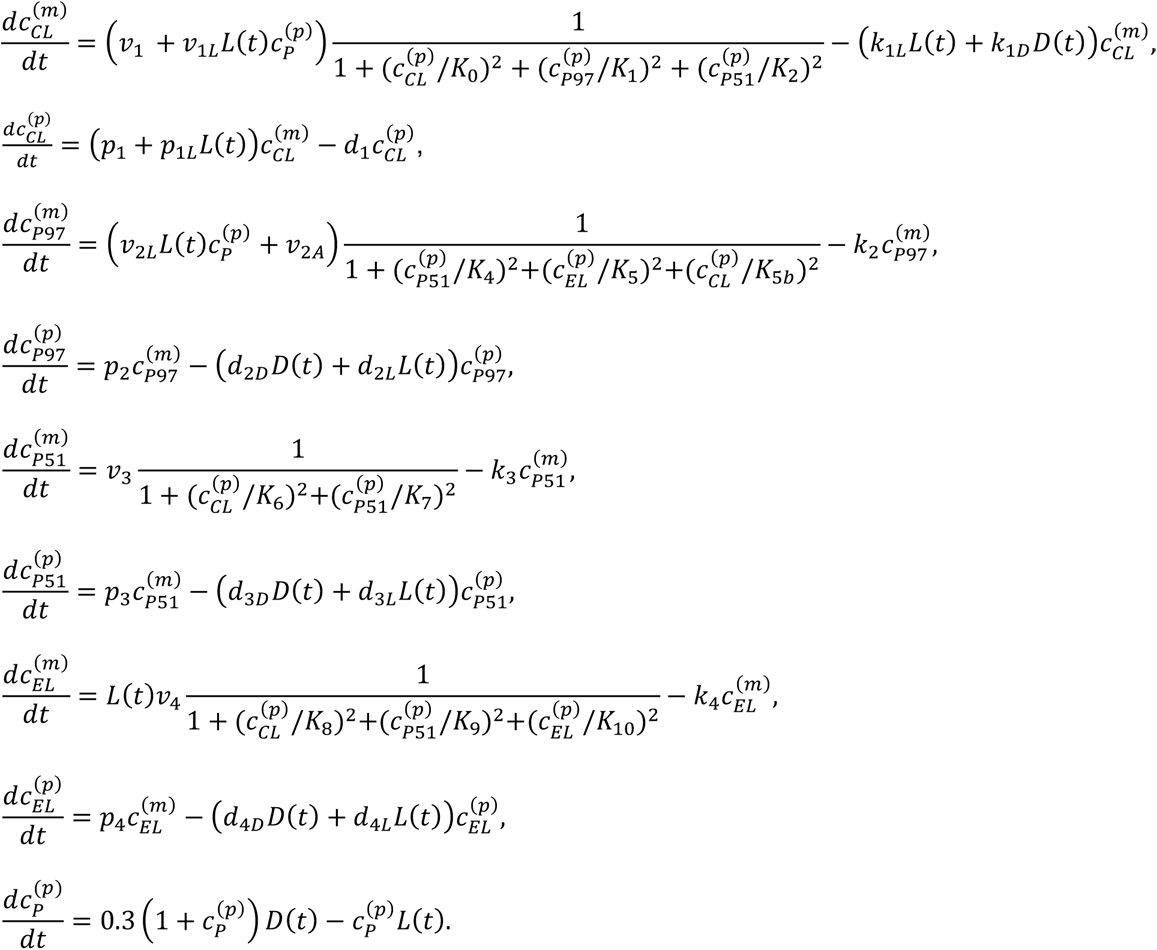

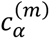 and 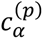 represent the concentration of the α-th mRNA and protein (or protein complex) respectively, for α = CL, P97, P51, EL, and P. *L*(t) represents the input light signal (*L =* 0, lights off; *L* > 0, lights on) and *D(t*) denotes a corresponding darkness input signal (*D =* 1, lights off; *D =* 0, lights on). The model contains 34 parameters, the values of which have been obtained previously through automated optimization [31]. The parameter values for *K*_0_, *K*_4_, *K*_5_, *K*_5*b*_ related to the modified interaction from CL to P97 were optimized by minimizing the cost function detailed in the following subsection. Sobol search and simulated annealing were combined to optimize within the range *K*_i_ ɛ [0.1,10] for *i* = 0,4,5,5*b*. The optimized parameter values together with the original values determined by De Caluwe et al. [31] are listed in S1 Table. The free running periods of the revised model are 25.5 h (LL; L = 1) and 26.25 h (DD; D = 1). Under LD cycles, peak times (ZT) of the gene expressions were CL = 0.50 h, P97 = 8.25 h, P51 = 17.50 h, and EL = 11.75 h.

### Cost function

In the parameter optimization of the single cell model, the cost function was computed as follows. First, the model was simulated under 12-h light–12-h dark conditions for a total of 60 days and then released into LL for 60 days, followed by constant dark (DD) for 60 days. In each light condition, the first 55 days were discarded as a transient dynamic. To ensure detectable rhythmicity under LL and DD conditions, all variables were required to have a minimum value of 0.1, as well as a minimum difference of 10 % between their minimum and maximum values. Any solution that did not meet these criteria was penalized with an arbitrarily large score. Then, the free-running period was calculated using the chi-square periodogram [47] from CL gene expression, at a significance level of 1 %. A score of 0 was given to a solution having a free-running period between 24 and 25 h under LL and between 25 and 28 h under DD. Solutions with free-running periods outside these ranges were allocated the scores of (*τ_LL_* − 24.5)^2^/(0.1·24.5)^2^ and (*τ_DD_* − 26.5)^2^/(0.1·26.5)^2^, where *τ_LL_* and *τ_DD_* represent free-running periods under LL and DD, respectively. For simulations under 12-h light–12-h dark cycles, solutions that were not entrained to the LD cycles were penalized with an arbitrarily large score. Entrained solutions were given a score of 0 for each gene that attained peak expression within ± 1 h of the expected ZT, which were as follows: CL = 1.5, P97 = 6, P51 = 12, and EL = 9. Expression peaks lying outside these intervals were scored as (*ZT_CL_* − 1.5)^2^/(0.1·24)^2^, (*ZT_P_*_97_ − 6)^2^/(0.1·24)^2^, (*ZT_P_*_51_ − 12)^2^/(0.1·24)^2^, (*ZT_EL_* − 9)^2^/(0.1·24)^2^, where *ZT*_α_ denotes Zeitgeber time of the α-th gene’s peak expression.

### Spatial molecular model

To simulate the spatial dynamics, we implemented the revised model on a simplified template of a plant. The template consisted of approximately 800 cells, classified into cotyledon, hypocotyl, root, and root tip regions (Fig 1B). Each cell contained an implementation of the revised compact model. To simulate growth of the seedlings, we added a row of cells to the root tip every 24 h. During this growth, the root tip region of the template was kept fixed in size. To do this, after the addition of new cells, the previously uppermost root tip cells became root cells instead.

As the coupling agents have yet to be clearly identified by experimental studies, the individual cells are assumed to be coupled through CL. Our model for coupled plant cells is described as follows:

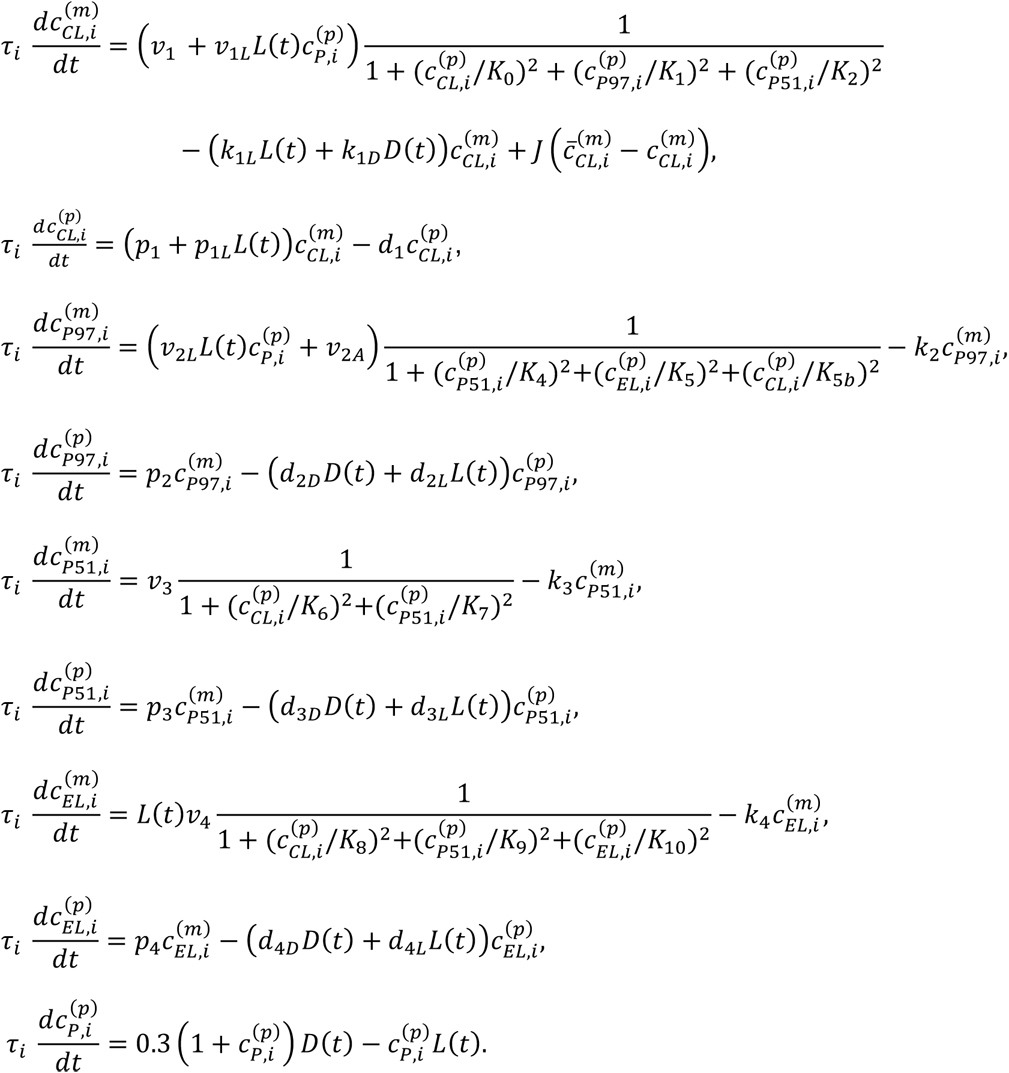

As in the revised single cell model, 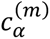 and 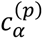 represent the concentration of the α-th mRNA and protein (or protein complex) respectively for α = CL, P97, P51, EL, and P, in the *i*-th cell (*i* = 1, 2,…, *N*). In the first equation, the CL gene is locally and diffusively coupled to its neighboring cells, where *J* is the coupling strength and 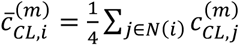 represents averaged expression level over 4 neighboring cells (left, right, above, and below). In the case of coupling between 8 neighboring cells (left, right, above, below, left above, left below, right above, and right below), the averaged expression level becomes 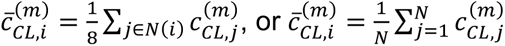 for the case of globally coupled cells. The coupling strength was set to *J* = 0, 0.001, 0.01, 0.1, 1, 2, or 4.

### Differential light input to cells

We set the light strength, *L*, to vary depending on the position of the cell. In doing so, we found that modest differences in *L* were sufficient to generate period differences between regions. To approximate the period differences that we observed between regions in experiments (Fig 2A) we set *L* = 1.35 for cotyledon cells, *L* = 1.10 for hypocotyl cells, *L* = 0.90 for root cells, and *L* = 1.03 for root tip cells. As the template of the seedling grew, we kept the root tip region of fixed size (see ‘Spatial molecular model’). This meant that the uppermost root tip cells become root cells instead. After this transition, the light input decreased so that cells received a level characteristic of root cells, *L* = 0.90. This caused these cells to slow.

To generate cell-to-cell variability in periods, the time scaling parameters *τ_i_* were set according to normal distributions as N(1, 0.059), N(1, 0.028), N(1, 0.073), N(1, 0.089) for the four regions, as informed by the analysis of single cell data [8]. For these values of *L* and *τ_i_*, the intrinsic periods of the 4 regions (*J* = 0) in the absence of growth were 23.2 ± 1.4 h, 25.5 ± 0.7 h, 27.9 ± 2.0 h, and 26.3 ± 2.3 h (mean ± standard deviation) respectively.

### Characterizing clock periods from experiments

We analyzed experimental luciferase and confocal data to characterize the periods within a seedling. We previously completed an organ-level analysis of period and phase for *PRR9*::*LUC*, *ELF4*::*LUC*, and *TOC1*::*LUC* expression [14]. In this analysis, 315 µm diameter regions of interest (ROI) were defined to represent the cotyledon, hypocotyl, root, and root tip regions of the seedling. These ROI were used to generate time series, from which period estimates were made across a number of individual seedlings. Here, we pooled estimates made from *PRR9*::*LUC*, *ELF4*::*LUC*, and *TOC1*::*LUC* lines, and calculated the mean for each region (Fig 2A). Only periods from time series classified as rhythmic (as defined previously [14]) were used in the calculation.

To characterize the variability of the periods, we measured the between-cell variation of clock periods in each region of the plant (S1 Fig). To do this we analyzed a dataset of single cell time-lapse microscopy, CCA1-YFP translational fusion data [8]. We defined cells as being within the root tip region if they are less than 315 µm from the most distal cell of the root tip. Where there are multiple imaging sections in the same organ, cells are pooled to give one region for the cotyledon, hypocotyl, and root. Variability was estimated using the coefficient of variation (CV; standard deviation / mean) of all cells within these regions. Only periods from time series classified as rhythmic (as defined previously [8]) were used in the calculation.

### Organ-level analysis

To assess our model we quantified the times of the peaks of expression. For experimental data, we used the ROI positions defined previously [14]. For the model, we defined 5-by-3 (width-by-height) pixel ROI on the seedling template, approximating the positions used in the experimental analysis of (S6 Fig). The mean of these ROI at each time point were taken to give the simulated time series. Peaks within 24–144 h after transfer to LL were identified using the “findpeaks” MATLAB (MathWorks, UK) function, with the constraint that peaks must be more than 19 h apart. For visual clarity, only peaks in which all organs completed the full cycle within the time window were plotted. Peaks of simulated expression were plotted against previously published experimental data for comparison (e.g. Fig 2D–F) [14]. Only peaks from experimental time series classified as rhythmic (as defined previously [14]) were plotted.

### Space-time plots

To visualize the spatial clock dynamics, we created space-time plots of gene expression (e.g. Fig 3A). We made plots of *PRR9*::*LUC*, *ELF4*::*LUC*, and *TOC1*::*LUC* expression as described previously [14]. Individual time series of the space-time plots were background subtracted and small gaps caused by segmentation errors were interpolated. A third-order Butterworth filter was then applied to reduce the high frequency noise. To create space-time plots of simulated clock gene expression, we also extracted the expression from sections that are perpendicular to the primary axis of the seedling (S6 Fig). We take the mean expression from 1-by-5 cell cross-sections of the template. To aid interpretation of space-time plots, amplitude trends were removed from both experimental and model space-time plots using part of the mFourFit algorithm [25].

We additionally plotted the times of the final peaks of expression of the space-time plots (e.g. Fig 3B). The final oscillation in which all organs completed the full cycle was used. We detected peaks within a 24 h window of the expected time of these peaks. If more than one peak was detected, we plotted the peak with the highest height.

### Entrainment to noisy LD cycles

We modelled entrainment by simulating light cycles with a day length of 12 h. The level of light during the day varied depending on the region of the template, matching *L* values under LL. We termed these cycles without noise ‘idealized LD’ (Fig 4A). To simulate fluctuations in the light cycle we utilized an algorithm developed previously [48], which approximates meteorological data [49]. Fluctuations lasted a random interval of time drawn from an exponential distribution of mean 2.4 h. The intensity of light during each fluctuation was drawn from a second uniform distribution of the range 0 to *L_max_*, with *L_max_* matching the *L* values for the cell under LL. The values are set according to the position of the cell on the template, as under LL. We termed this condition ‘noisy LD’ (Fig 4D). We made the simplifying assumption that each cell is exposed to an independent noisy LD cycle due to their unique positions in the environment. LD cycles were input to the molecular model through the parameter *L*.

### Synchrony analysis

We estimated the global synchrony in simulations from the variance of the individual cells (Fig 5 and S14 Fig). Firstly, the average signal of all *N* cells of the simulation {*x_h_*(*t*): *h* = 1, 2,…, *N*}, was computed as 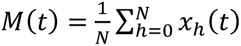. Then the overall level of synchrony was measured by the synchronization index [38],

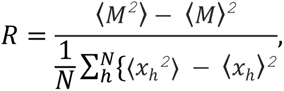

where <> denotes the time average, which in our case was between 0 and 144 h. The synchronization index measures the ratio of the variance of the average signal to the average variance of the individual cells. The parameter quantifies the distribution of both the phases and amplitudes of the individual cells. However, we normalized the individual traces to remove amplitude variance and therefore measured only the phase differences. *R* ranges from 1 when cells are synchronized, to 0 when cells are completely desynchronized.

**S1 Video. Time-lapse of simulated *PRR9*/*PRR7* expression under LL.** Representative simulation of simulated *PRR9*/*PRR7* expression from individual cells of the plant template 24– 144 h after transfer to LL. LL, constant light; *PRR*, *PSEUDO RESPONSE REGULATOR*.

**S2 Video. Time-lapse of *PRR9*::*LUC* expression under LL.** *PRR9*::*LUC* expression from a single representative seedling 24–144 h after transfer to LL. LL, constant light; *PRR9*, *PSEUDO RESPONSE REGULATOR9*.

---

**S1 Fig.**
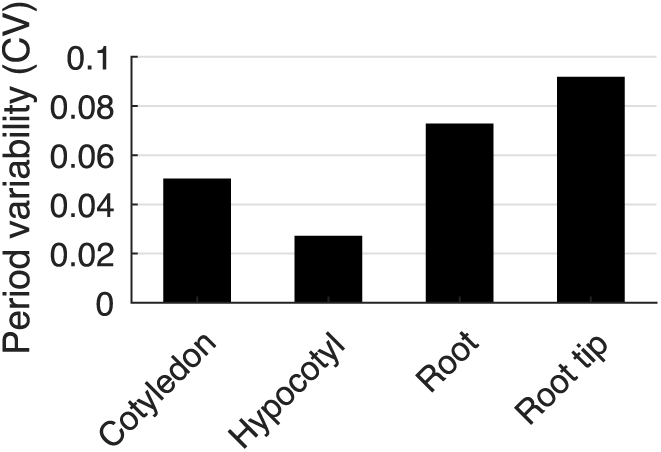
Between-cell variability of CCA1-YFP rhythms. Between-cell variability of periods within different organs. The periods were estimated from single cell time-lapse movies of CCA1- YFP expression carried out previously [8]. *CCA1*, *CIRCADIAN CLOCK ASSOCIATED 1*; CV, coefficient of variation; YFP, yellow fluorescent protein.

**S2 Fig.**
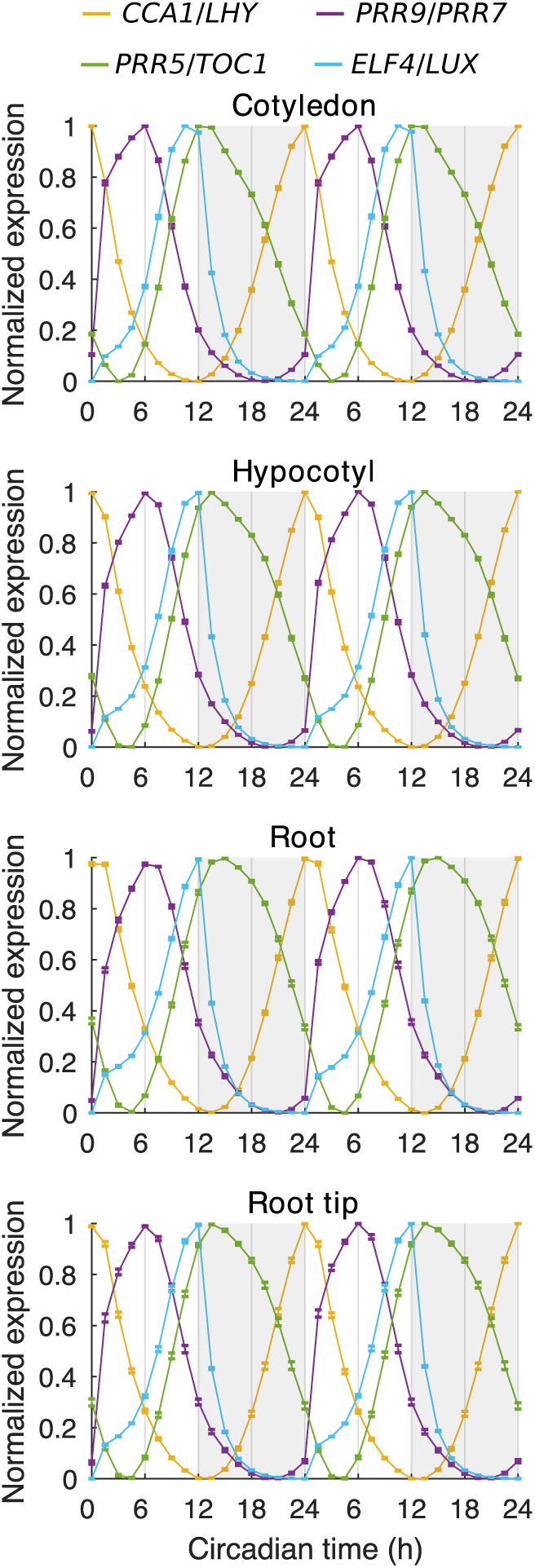
Simulated expression of clock genes in different regions under LD cycles. Expression of simulated *CCA1*/*LHY*, *PRR9*/*PRR7*, *PRR5*/*TOC1*, and *ELF4*/*LUX* from regions of the seedling template 96–144 h after the beginning of LD cycles. The light input, *L*, was varied depending on whether the cell was within the cotyledon (*L* = 1.35), hypocotyl (*L* = 1.10), root (*L* = 0.90), or root tip (*L* = 1.03) region, and coupling was present between cells (*J* = 2). Data represent the mean ± standard error, *n* = 9 simulations. *CCA1*, *CIRCADIAN CLOCK ASSOCIATED 1*; *ELF4*, *EARLY FLOWERING 4*; LD, light-dark; *LHY*, *LATE ELONGATED HYPOCOTYL; LUC*, *LUCIFERASE*; *LUX*, *LUX ARRHYTHMO*; *PRR*, *PSEUDO-RESPONSE REGULATOR*; *TOC1*, *TIMING OF CAB EXPRESSION 1*.

**S3 Fig.**
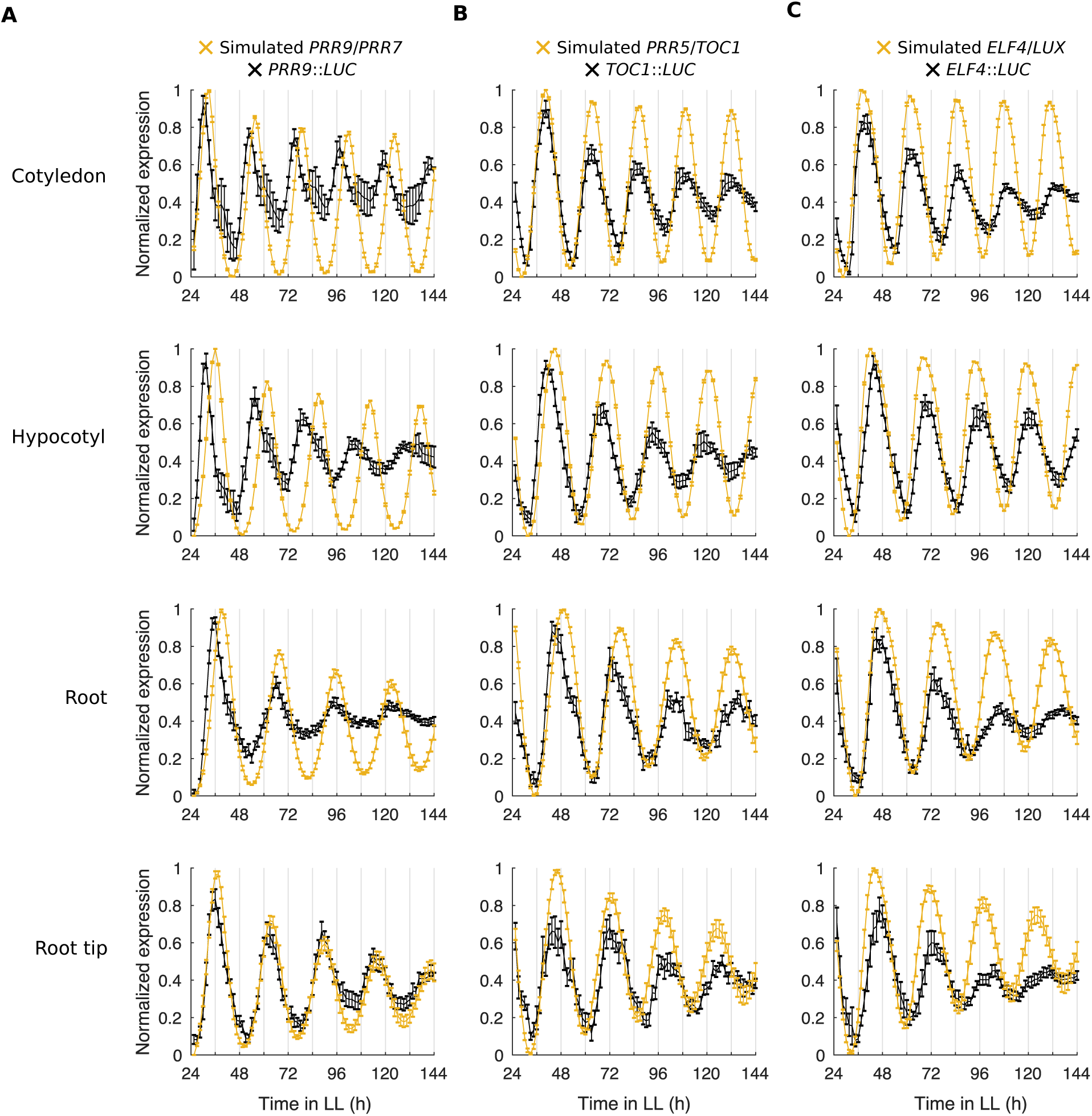
Simulated expression of multiple core clock genes under LL qualitatively match luciferase reporter experiments. (A–C) Expression of simulated *PRR9*/*PRR7* and *PRR9*::*LUC* (A), simulated *PRR5*/*TOC1* and *TOC1*::*LUC (B)*, or simulated *ELF4*/*LUX* and *ELF4*::*LUC* (C) in different organs under LL. Simulations include different levels of light input to regions and cell-to-cell coupling (*J* = 2). Data represent the mean ± standard error of organs scored as rhythmic. *n* = 9 simulations. Experimental data is from time-lapse movies carried out previously [14]. *ELF4*, *EARLY FLOWERING 4*; LL, constant light; *LUC*, *LUCIFERASE*; *LUX*, *LUX ARRHYTHMO*; *PRR*, *PSEUDO-RESPONSE REGULATOR*; *TOC1*, *TIMING OF CAB EXPRESSION 1*.

**S4 Fig.**
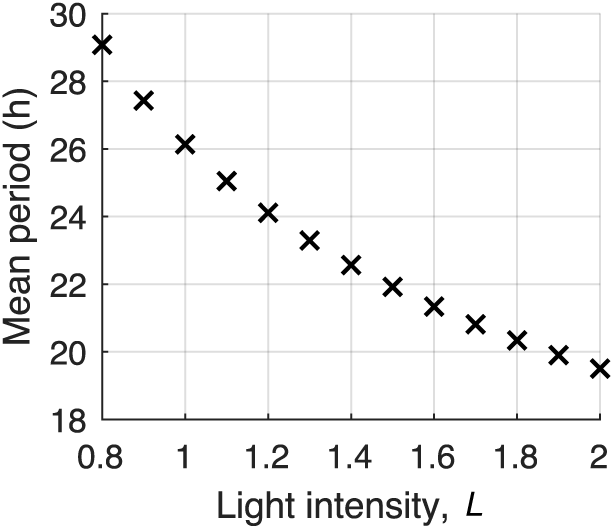
Higher light intensities cause simulations to run faster. The mean peak-to-peak time of simulated *PRR9*/PRR*7* expression with increasing intensities of light, *L*. A single instance of the model was implemented at each *L*, without cell-to-cell coupling or variation in gene expression. *L* values below 0.8 caused damping in the original and modified De Caluwe implementation. *PRR*, *PSEUDO-RESPONSE REGULATOR*.

**S5 Fig.**
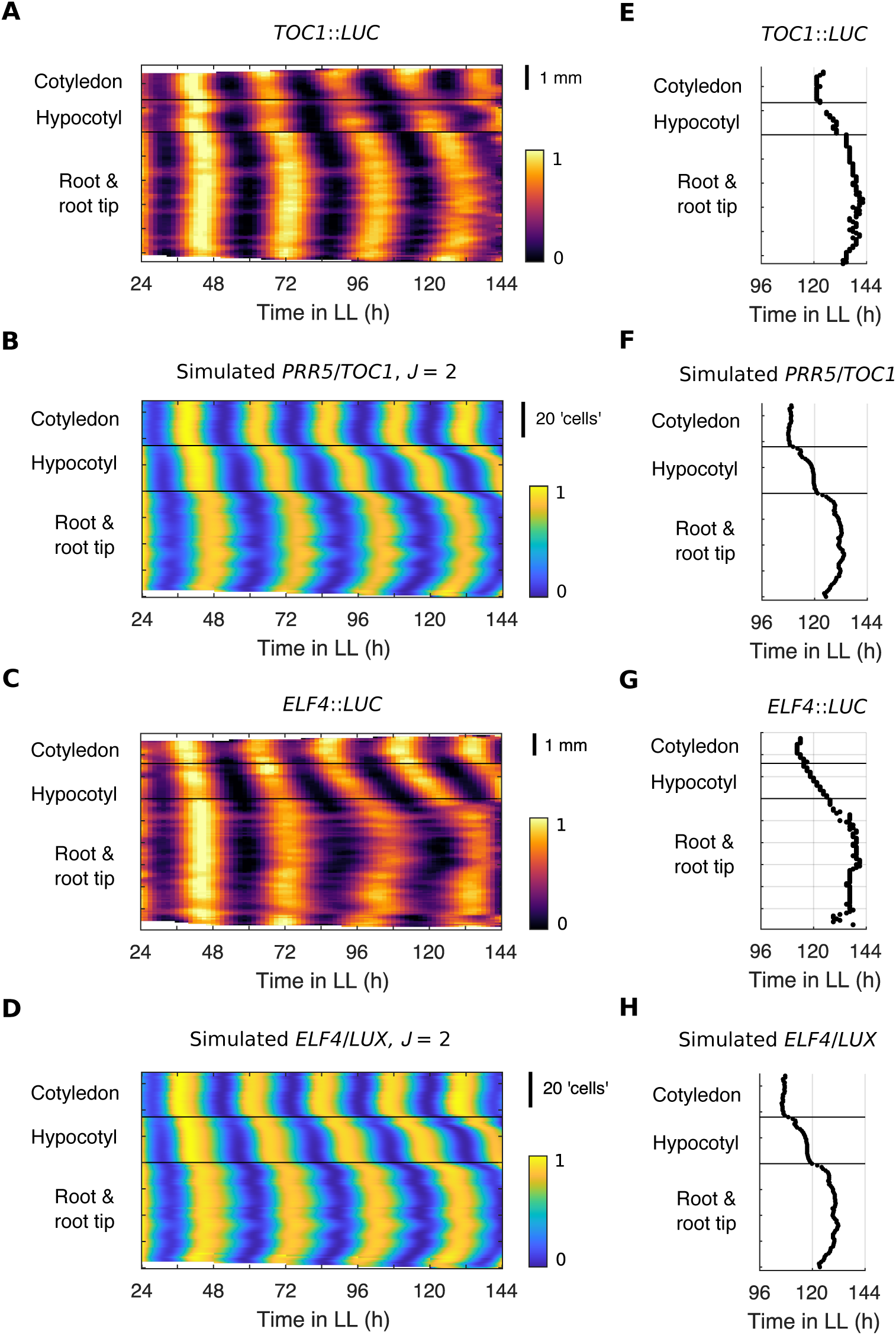
Local sharing of mRNA can reproduce experimentally observed spatial waves of clock gene expression for multiple core clock genes. (A–D) Representative intensity plot of *TOC1*::*LUC* (A), simulated *PRR5*/*TOC1* (B)*, ELF4*::*LUC* (C) and simulated *ELF4*/*LUX* (D) expression across longitudinal sections of a single seedling under LL. Simulations include different levels of light input to regions and cell-to-cell coupling (*J* = 2). (E–H) Times of the final peak of *TOC1*::*LUC* (E), simulated *PRR5*/*TOC1* (F)*, ELF4*::*LUC* (G) and simulated *ELF4*/*LUX* (H) intensity plots. Experimental data is an analysis of time-lapse movies carried out previously [14]. *ELF4*, *EARLY FLOWERING 4*; *LHY*, *LATE ELONGATED HYPOCOTYL*; LL, constant light; *LUX*, *LUX ARRHYTHMO*; *PRR5*, *PSEUDO-RESPONSE REGULATOR 5; TOC1*, *TIMING OF CAB EXPRESSION 1*.

**S6 Fig.**
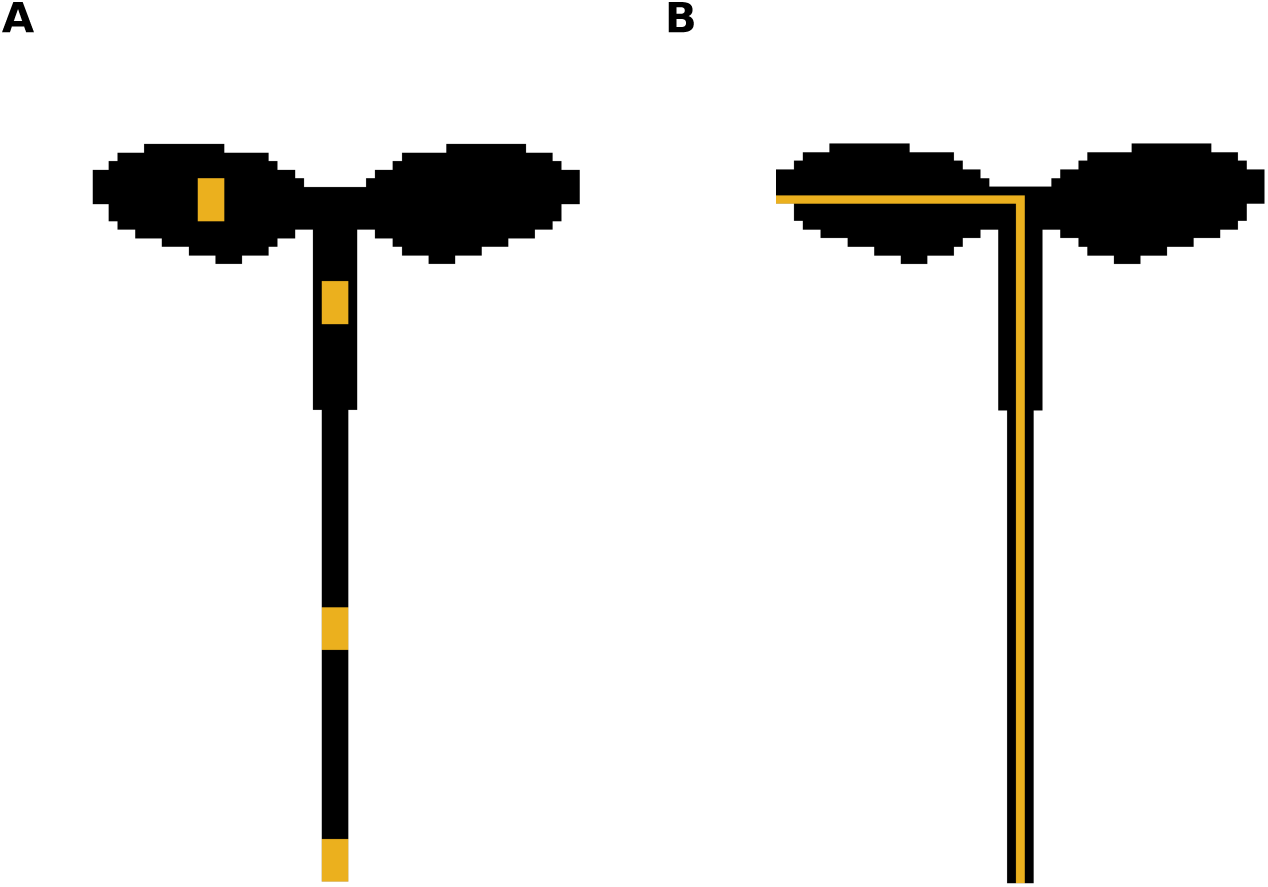
Spatial analysis of rhythms across the simplified template of a plant. (A) To measure the period and phase of an organ, the expression was extracted from 3-by-3 pixel regions of interest (yellow squares). (B) To produce space-time plots of gene expression (e.g. Fig 3A), the mean expression of sections perpendicular across the central axis (yellow line) of the template were taken. Luciferase space-time plots were created similarly, as described previously [14] (Methods).

**S7 Fig.**
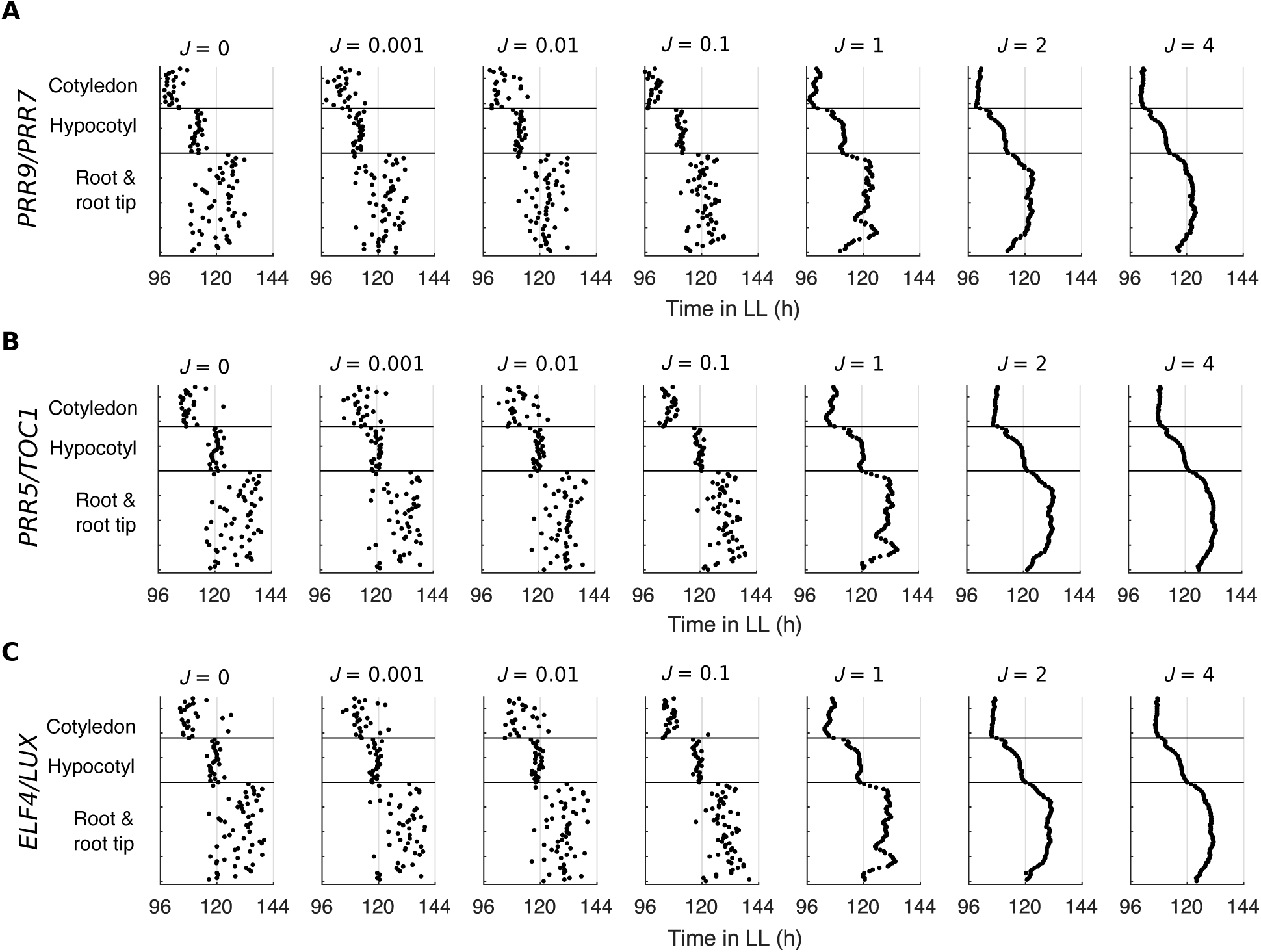
Times of the final peaks of simulated expression under LL with increasing strengths of coupling. (A–C) Times of the final peak of simulated *PRR9/PRR7* (A), simulated *PRR5*/*TOC1* (B), and simulated *ELF4*/*LUX* (C) intensity plots, each simulated under LL with increasing strength of coupling, *J*. *ELF4*, *EARLY FLOWERING 4*; LL, constant light; *LUX*, *LUX ARRHYTHMO*; *PRR*, *PSEUDO-RESPONSE REGULATOR; TOC1*, *TIMING OF CAB EXPRESSION 1*.

**S8 Fig.**
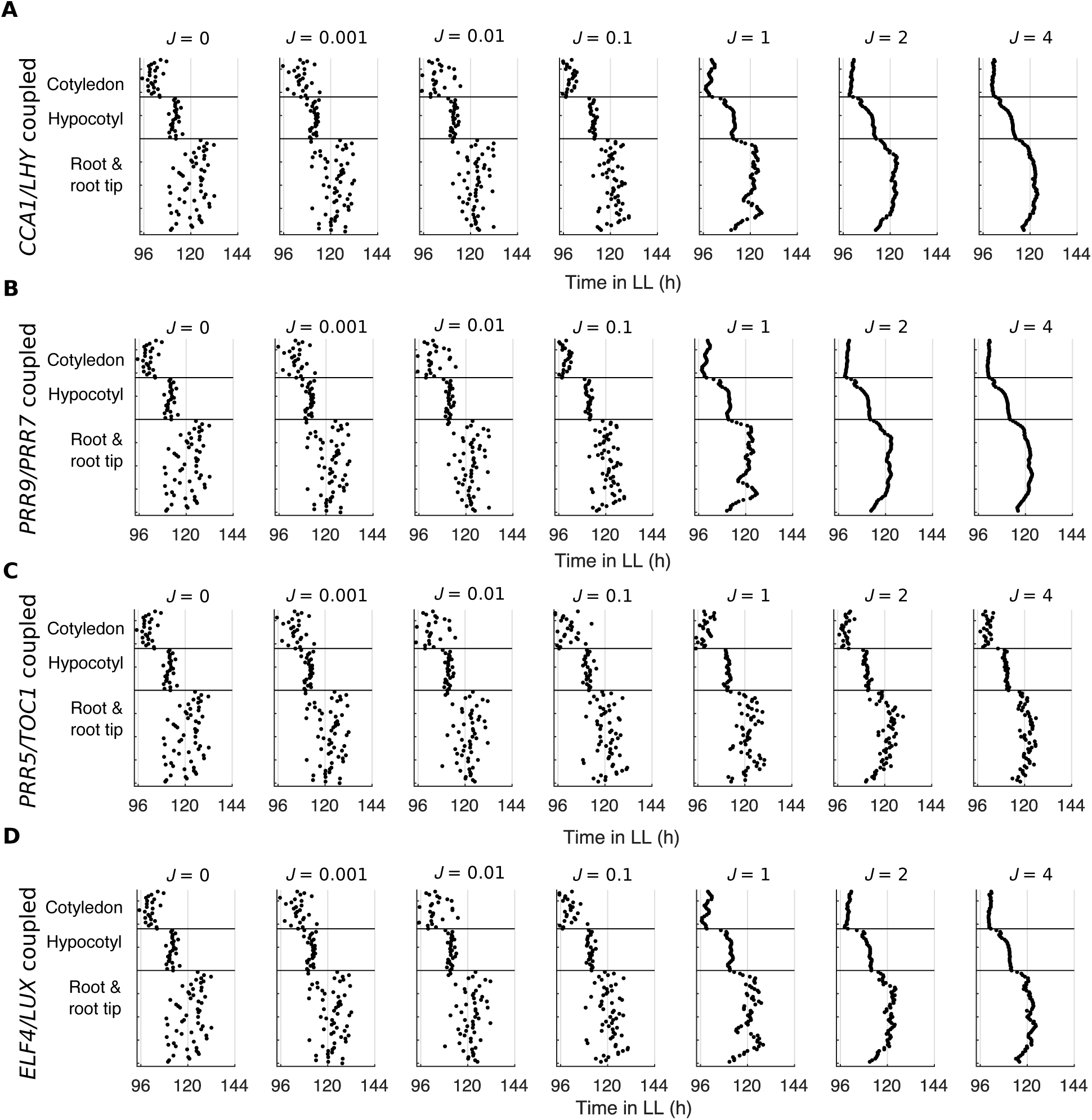
Times of the final peaks of simulated expression under LL with the sharing of different clock gene mRNA between cells. (A–C) Times of the final peak of simulated *PRR9/PRR7* intensity plots, each simulated with the sharing of *CCA1*/*LHY*, *PRR9*/*PRR7*, *PRR5*/*TOC1*, or *ELF4*/*LUX* mRNA between neighbor cells. Simulations were under LL with increasing strength of coupling, *J*. *CCA1*, *CIRCADIAN CLOCK ASSOCIATED 1*; *ELF4*, *EARLY FLOWERING 4*; *LHY*, *LATE ELONGATED HYPOCOTYL*; LL, constant light; *LUX*, *LUX ARRHYTHMO*; *PRR*, *PSEUDO-RESPONSE REGULATOR; TOC1*, *TIMING OF CAB EXPRESSION 1*.

**S9 Fig.**
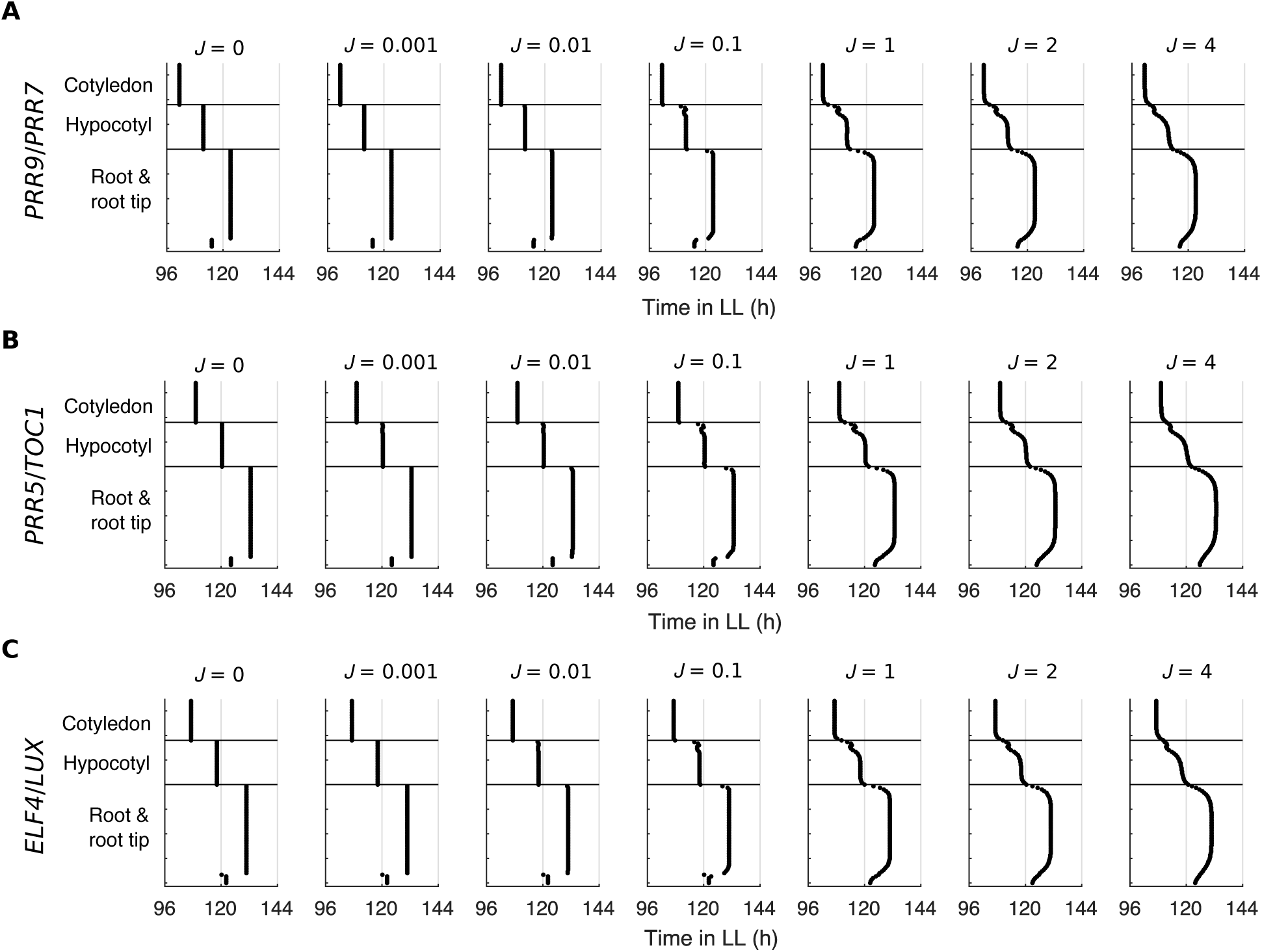
Times of the final peaks of simulated expression under LL without variation in gene expression. (A–C) Times of the final peak of simulated *PRR9*/*PRR7* (A)*, PRR5*/*TOC1* (B), or *ELF4*/*LUX* (C) intensity plots, each simulated with the sharing of *CCA1*/*LHY* mRNA between 4 neighbor cells, but without cell-to-cell variation in gene expression. Simulations were under LL with increasing strength of coupling, *J*. *CCA1*, *CIRCADIAN CLOCK ASSOCIATED 1*; *ELF4*, *EARLY FLOWERING 4*; *LHY*, *LATE ELONGATED HYPOCOTYL*; LL, constant light; *LUX*, *LUX ARRHYTHMO*; *PRR*, *PSEUDO-RESPONSE REGULATOR; TOC1*, *TIMING OF CAB EXPRESSION 1*.

**S10 Fig.**
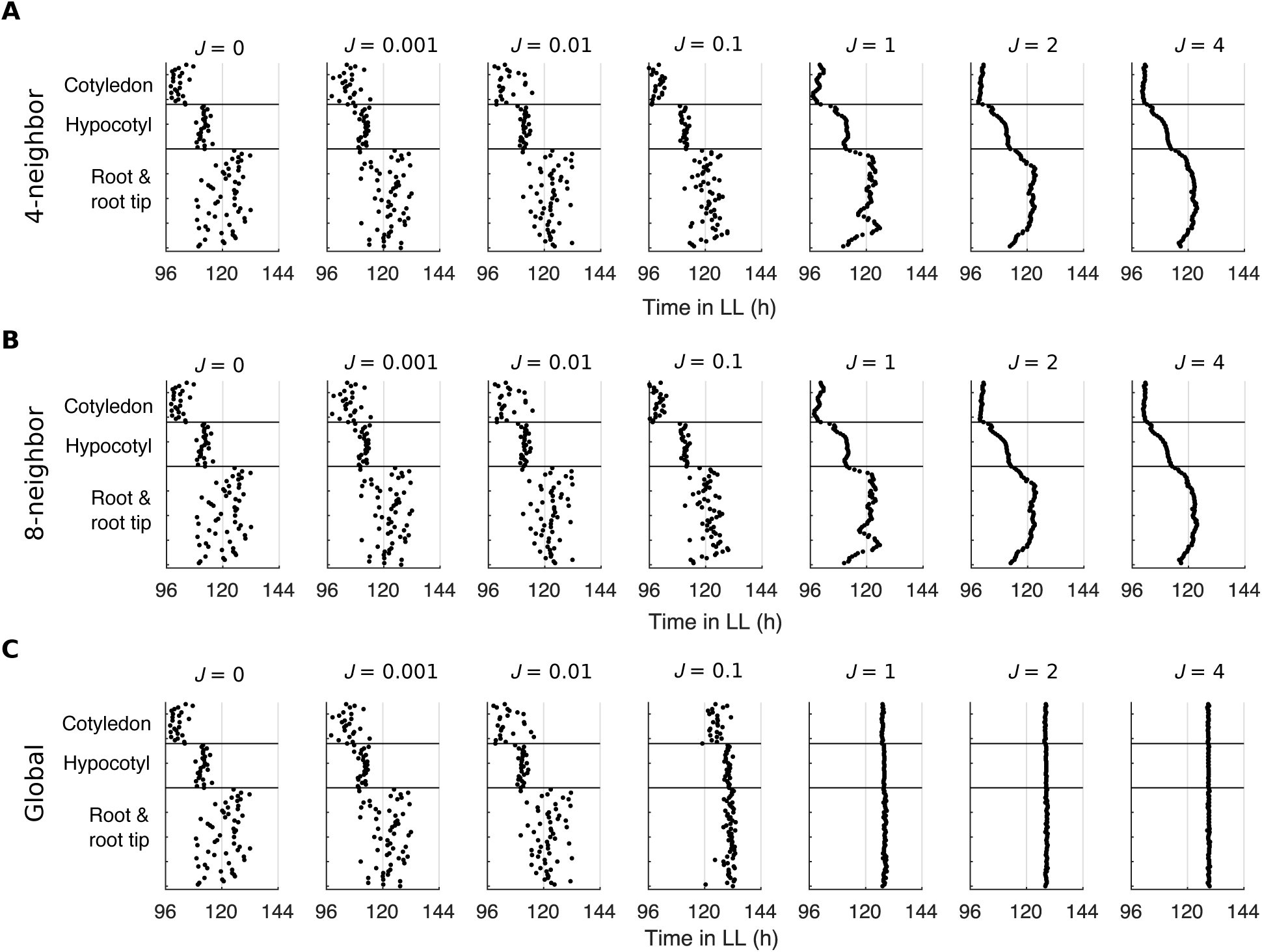
Times of the final peaks of simulated expression under LL with different coupling rules. (A–C) Times of the final peak of simulated *PRR9/PRR7* intensity plots, each simulated under LL with the sharing of *CCA1*/*LHY* mRNA between the 4 nearest neighbor cells (A), 8 nearest neighbor cells (B), or globally (all-to-all) (C). Simulations were run under LL with increasing strength of coupling, *J*. *CCA1*, *CIRCADIAN CLOCK ASSOCIATED 1*; *LHY*, *LATE ELONGATED HYPOCOTYL*; LL, constant light; *PRR*, *PSEUDO-RESPONSE REGULATOR*.

**S11 Fig.**
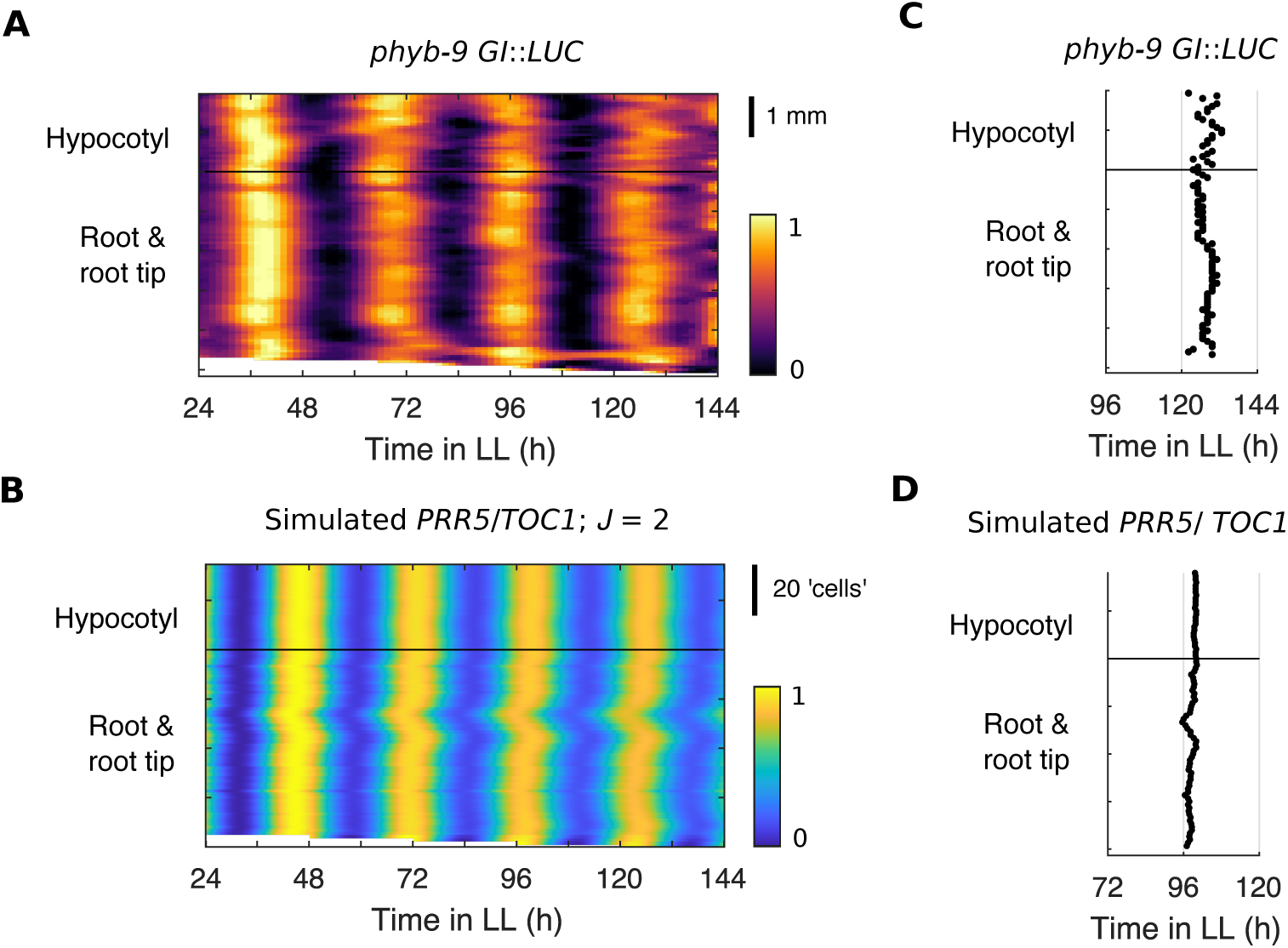
Simulations with equal light input to regions predicts the loss of spatial waves observed in the *phyb-9* mutant. (A) Representative intensity plot of *GI*::*LUC* expression across longitudinal sections of a single seedling under constant red light. Imaging was performed in the light sensing mutant *phyb-9* background. (B) Representative intensity plot of simulated *PRR5*/*TOC1* expression across longitudinal sections of a single seedling under LL. Simulations were performed with equal low light to all regions (*L* = 0.85) and cell-to-cell coupling (*J* = 2). (C) Times of the final peaks of *phyb-9 TOC1*::*LUC* intensity plots. (D) Times of the final peaks of simulated *PRR5*/*TOC1* intensity plots, with equal low light to all regions (*L* = 0.85) and cell-to-cell coupling (*J* = 2). Experimental data is an analysis of time-lapse movies carried out previously [14]. Intensity plots are limited to the lower hypocotyl and root due to rapid elongation of the hypocotyl in the *phyb-9* mutant. *PRR*, *PSEUDO RESPONSE REGULATOR*; LL, constant light; *LUC*, *LUCIFERASE*. *TOC1, TIMING OF CAB EXPRESSION 1*.

**S12 Fig.**
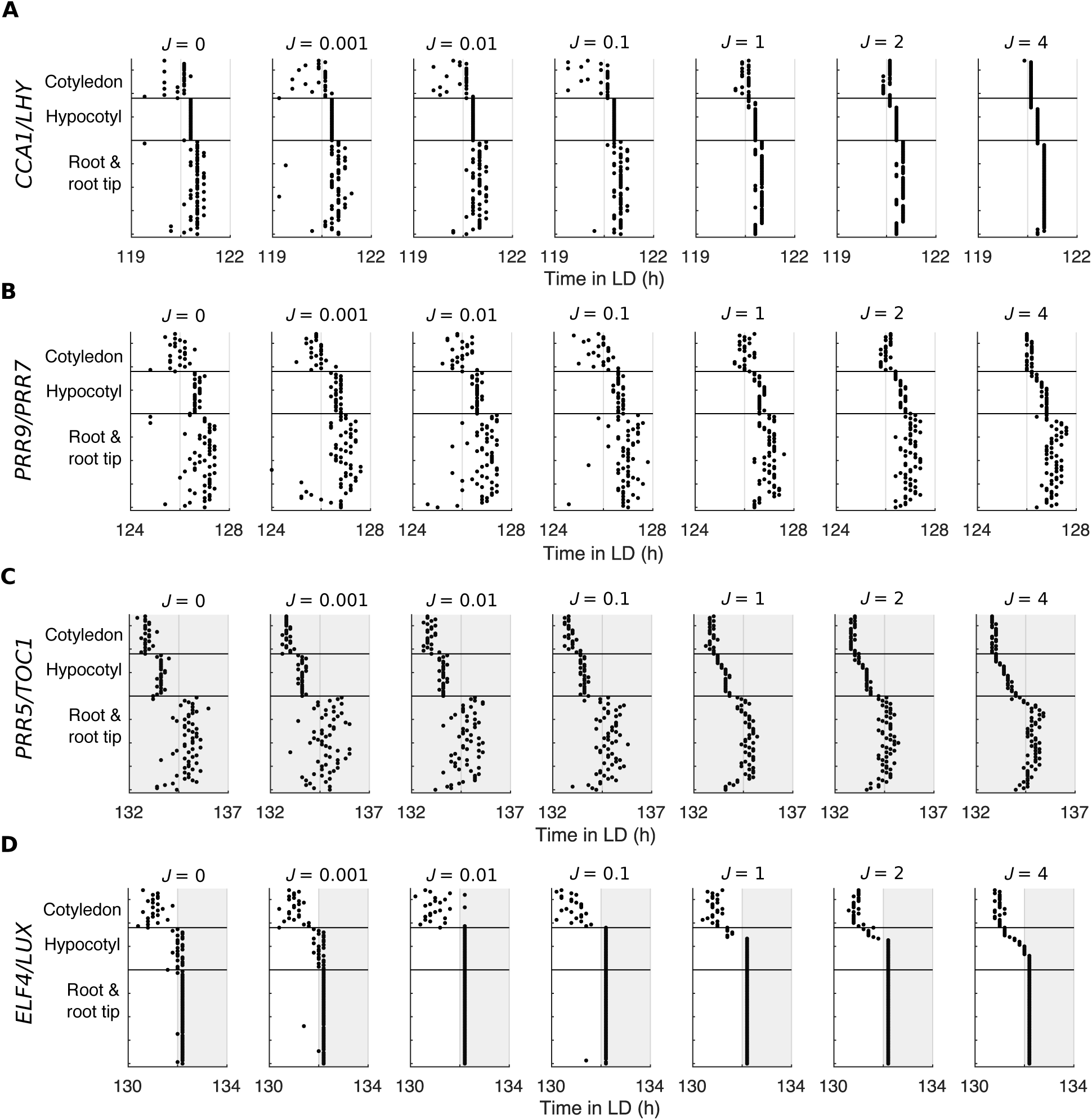
Times of the final simulated peaks of expression under idealized LD cycles with increasing strengths of coupling. (A–D) Times of the final peak of simulated *CCA1*/*LHY* (A), *PRR9/PRR7* (B), *PRR5*/*TOC1* (C), and *ELF4*/*LUX* (D) intensity plots, each simulated under idealized LD cycles with increasing strength of coupling, *J*. Cells were exposed to idealized LD cycles during simulations. Grey background indicates nighttime. *ELF4* appears synchronized when the peak coincides with the dark transition as darkness causes strong repression of *ELF4* expression. *CCA1*, *CIRCADIAN CLOCK ASSOCIATED 1*; *ELF4*, *EARLY FLOWERING 4*; LD, light-dark; *LHY*, *LATE ELONGATED HYPOCOTYL*; *LUX*, *LUX ARRHYTHMO*; *PRR*, *PSEUDO RESPONSE REGULATOR*; *TOC1*, *TIMING OF CAB EXPRESSION 1*.

**S13 Fig.**
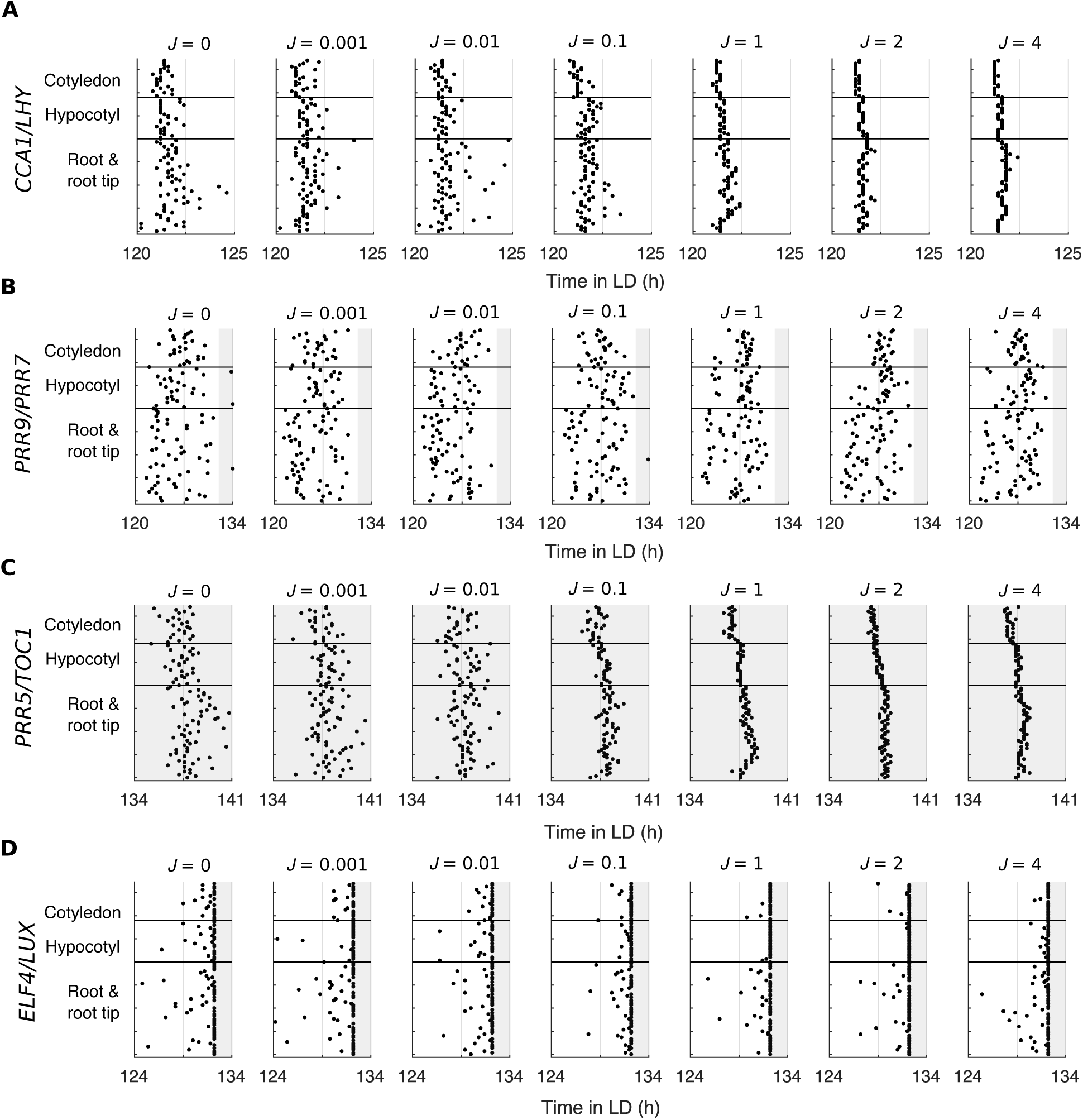
Times of the final simulated peaks of expression under noisy LD cycles with increasing strengths of coupling. (A–D) Times of the final peak of simulated *CCA1*/*LHY* (A), *PRR9/PRR7* (B), simulated *PRR5*/*TOC1* (C), and simulated *ELF4*/*LUX* (D) intensity plots, each simulated under idealized LD cycles with increasing strength of coupling, *J*. Cells were exposed to idealized LD cycles during simulations. Grey background indicates nighttime. *ELF4* appears synchronized when the peak coincides with the dark transition as darkness causes strong repression of *ELF4* expression. *CCA1*, *CIRCADIAN CLOCK ASSOCIATED 1*; *ELF4*, *EARLY FLOWERING 4*; LD, light-dark; *LHY*, *LATE ELONGATED HYPOCOTYL*; *LUX*, *LUX ARRHYTHMO*; *PRR*, *PSEUDO RESPONSE REGULATOR*; *TOC1*, *TIMING OF CAB EXPRESSION 1*.

**S14 Fig.**
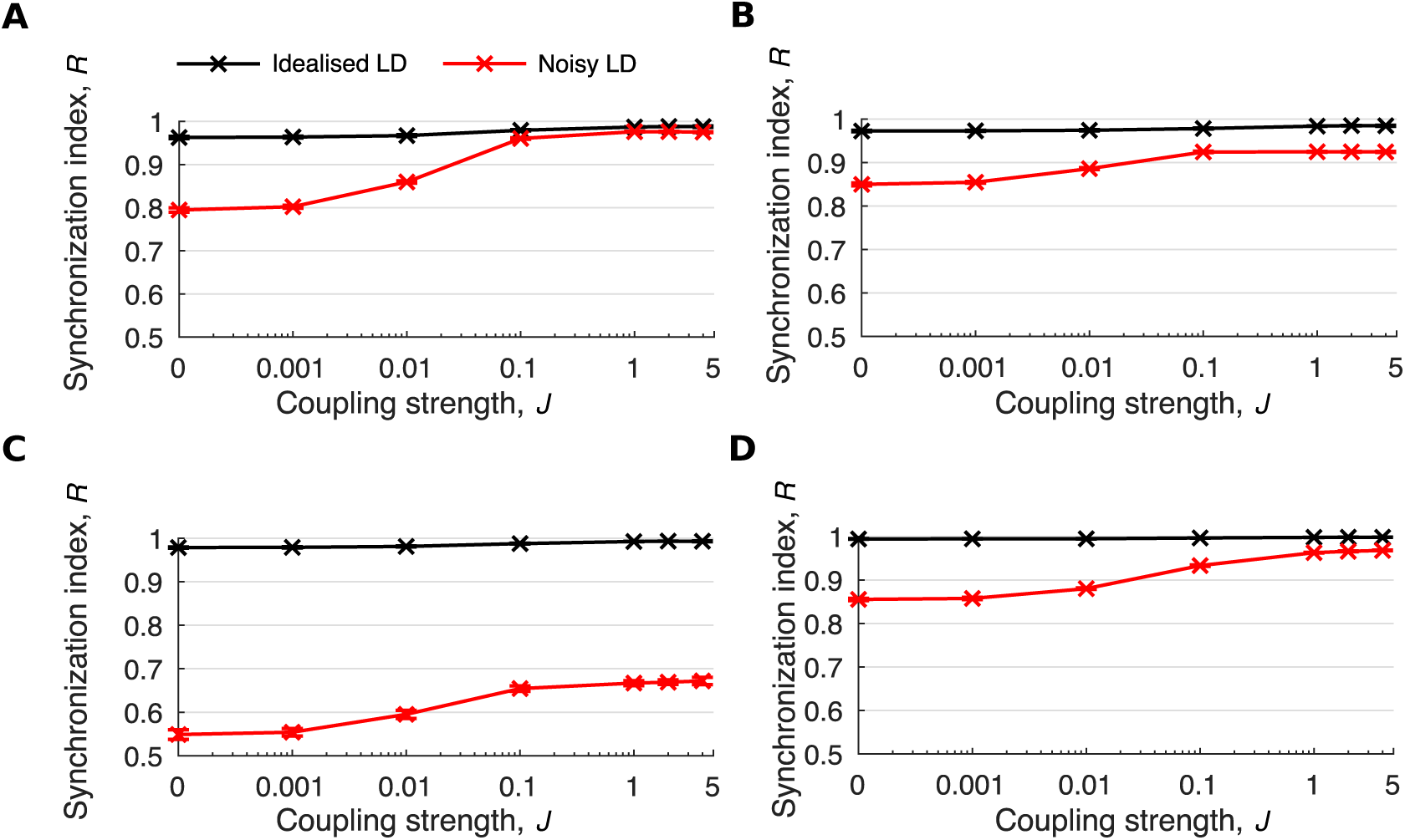
Cell-to-cell coupling increases the synchrony of protein oscillations under realistic LD cycles. (A–D) Quantification of phase coherence by the synchronization index, *R*, for simulated *CCA1*/*LHY* (A), *PRR9*/*PRR7* (B), *PRR5*/*TOC1* (C) or *ELF4*/*LUX* (D) protein expression under idealized or noisy LD cycles. Simulations included increasing strengths of coupling, *J*. Color legends are as in A. Data points represent the mean ± standard deviation, *n* = 9 simulations. CCA1, CIRCADIAN CLOCK ASSOCIATED 1; ELF4, EARLY FLOWERING 4; LD, light-dark; LHY, LATE ELONGATED HYPOCOTYL; LUX, LUX ARRHYTHMO; PRR, PSEUDO RESPONSE REGULATOR; TOC1, TIMING OF CAB EXPRESSION 1.

**S1 Table.** Parameter values for the spatial clock model.

| Parameter | Description | Original value<br>[31] | Re-<br>optimized<br>value |
| --- | --- | --- | --- |
| $v_1$ | *CCA1/LHY* synthesis | 4.6 nM h<sup>-1</sup> | 4.58 nM h<sup>-1</sup> |
| $v_{1L}$ | *CCA1/LHY* light-induced synthesis | 3.0 nM h<sup>-1</sup> | 3.0 nM h<sup>-1</sup> |
| $v_{2A}$ | *PRR9/PRR7* synthesis | 1.3 nM h<sup>-1</sup> | 1.27 nM h<sup>-1</sup> |
| $v_{2L}$ | *PRR9/PRR7* light-induced synthesis | 5.0 nM h<sup>-1</sup> | 5.0 nM h<sup>-1</sup> |
| $v_3$ | *PRR5/TOC1* synthesis | 1.0 nM h<sup>-1</sup> | 1.0 nM h<sup>-1</sup> |
| $v_4$ | *ELF4/LUX* synthesis | 1.5 nM h<sup>-1</sup> | 1.47 nM h<sup>-1</sup> |
| $k_{1L}$ | *CCA1/LHY* mRNA degradation (light) | 0.53 h<sup>-1</sup> | 0.53 h<sup>-1</sup> |
| $k_{1D}$ | *CCA1/LHY* mRNA degradation (dark) | 0.21 h<sup>-1</sup> | 0.21 h<sup>-1</sup> |
| $k_2$ | *PRR9/PRR7* mRNA degradation | 0.35 h<sup>-1</sup> | 0.35 h<sup>-1</sup> |
| $k_3$ | *PRR5/TOC1* mRNA degradation | 0.56 h<sup>-1</sup> | 0.56 h<sup>-1</sup> |
| $k_4$ | *ELF4/LUX* mRNA degradation | 0.57 h<sup>-1</sup> | 0.57 h<sup>-1</sup> |
| $p_1$ | *CCA1/LHY* translation | 0.76 h<sup>-1</sup> | 0.76 h<sup>-1</sup> |
| $p_{1L}$ | *CCA1/LHY* light-induced translation | 0.42 h<sup>-1</sup> | 0.42 h<sup>-1</sup> |
| $p_2$ | *PRR9/PRR7* translation | 1.0 h<sup>-1</sup> | 1.01 h<sup>-1</sup> |
| $p_3$ | *PRR5/TOC1* translation | 0.64 h<sup>-1</sup> | 0.64 h<sup>-1</sup> |
| $p_4$ | *ELF4/LUX* translation | 1.0 h<sup>-1</sup> | 1.01 h<sup>-1</sup> |
| $d_1$ | *CCA1/LHY* degradation | 0.68 h<sup>-1</sup> | 0.68 h<sup>-1</sup> |
| $d_{2D}$ | *PRR9/PRR7* degradation (dark) | 0.51 h<sup>-1</sup> | 0.5 h<sup>-1</sup> |
| $d_{2L}$ | *PRR9/PRR7* degradation (light) | 0.30 h<sup>-1</sup> | 0.29 h<sup>-1</sup> |
| $d_{3D}$ | *PRR5/TOC1* degradation (dark) | 0.48 h<sup>-1</sup> | 0.48 h<sup>-1</sup> |
| $d_{3L}$ | *PRR5/TOC1* degradation (light) | 0.78 h<sup>-1</sup> | 0.78 h<sup>-1</sup> |
| $d_{4D}$ | *ELF4/LUX* degradation (dark) | 1.2 h<sup>-1</sup> | 1.21 h<sup>-1</sup> |
| $d_{4L}$ | *ELF4/LUX* degradation (light) | 0.38 h<sup>-1</sup> | 0.38 h<sup>-1</sup> |
| $K_0$ | Inhibition of *CCA1/LHY* by *CCA1/LHY* | n/a | 5.07 nM |
| $K_1$ | Inhibition of *CCA1/LHY* by *PRR9/PRR7* | 0.16 nM | 0.16 nM |
| $K_2$ | Inhibition of *CCA1/LHY* by *PRR5/TOC1* | 1.2 nM | 1.18 nM |
| $K_4$ | Inhibition of *PRR9/PRR7* by *PRR5/TOC1* | 0.23 nM | 0.40 nM |
| $K_5$ | Inhibition of *PRR9/PRR7* by *ELF4/LUX* | 0.30 | 0.62 nM |
| $K_{5b}$ | Inhibition of *PRR9/PRR7* by *CCA1/LHY* | n/a | 1.2 nM |
| $K_6$ | Inhibition of *PRR5/TOC1* by *CCA1/LHY* | 0.46 | 0.46 nM |
| $K_7$ | Inhibition of *PRR5/TOC1* by *PRR5/TOC1* | 2.0 | 2.0 nM |
| $K_8$ | Inhibition of *ELF4/LUX* by *CCA1/LHY* | 0.36 | 0.36 nM |
| $K_9$ | Inhibition of *ELF4/LUX* by *PRR5/TOC1* | 1.9 | 1.9 nM |
| $K_{10}$ | Inhibition of *ELF4/LUX* by *ELF4/LUX* | 1.9 | 1.9 nM |

*CCA1*, *CIRCADIAN CLOCK ASSOCIATED 1*; *ELF4*, *EARLY FLOWERING 4*; *LHY*, *LATE* *ELONGATED HYPOCOTYL*; *LUX*, *LUX ARRHYTHMO*; *PRR*, *PSEUDO-RESPONSE* *REGULATOR*; *TOC1*, *TIMING OF CAB EXPRESSION 1*.

